# Assessing *Corynebacterium glutamicum* as a surrogate of *Mycobacterium tuberculosis* for DNA gyrase inhibitor design

**DOI:** 10.64898/2026.06.24.734172

**Authors:** Yaëlle Wormser, Emilie Yab, Adrià Sogues, Francesca Gubellini, Estelle Capton, Estelle Lecat, Mathilde Ben Assaya, Alexandra Aubry, Ariel Mechaly, Pedro M. Alzari, Anne Marie Wehenkel, Antoine Gedeon, Stéphanie Petrella

## Abstract

DNA gyrase is an essential bacterial enzyme and a clinically validated target for the treatment of tuberculosis. However, the discovery of new inhibitors remains limited by the many challenges regarding the manipulation on pathogenic mycobacteria. This study validates *Corynebacterium glutamicum* (*Cglu*) as a safe, non-pathogenic surrogate for *Mycobacterium tuberculosis* (*Mtb*) to investigate DNA gyrase and facilitate the identification of new inhibitors. Using *Cglu* as a target allows for fast whole-cell screening under safe conditions while ensuring efficient drug uptake. *Cglu* shares key physiological features with *Mtb*, including genome size, complex cell wall structure, and a single type I and type II topoisomerase. Structural and functional comparisons emphasize the similarity of *Cglu* and *Mtb* gyrases, which share 70% sequence identity and show comparable catalytic properties and responsiveness to known inhibitors. Thus, the cryo-EM structure of the *Cglu* gyrase-DNA complex at 3.2 Å resolution reveals highly conserved drug-binding pockets for known anti-gyrase inhibitors and the genetic depletion of *gyrA* or *gyrB* in *Cglu* causes severe growth and morphological defects, mirroring the effects of chemical inhibition and allowing to link gyrase function to cellular phenotypes. Comparative imaging of different inhibitor classes (fluoroquinolones, aminocoumarins, NBTIs) uncovers distinct morphological signatures that reflect each compound’s mode of action. Finally, cross-species complementation confirms functional conservation but also highlights subtle structural differences affecting efficiency. Together, these findings establish *Cglu* as a robust and biosafe model for dissecting gyrase function, visualizing DNA topology dynamics, and accelerating the discovery of gyrase-targeting antimicrobials. More generally, our studies demonstrate the feasibility of using *Cglu* as a cell-based screening platform to discover new anti-tuberculous compounds targeting conserved mechanisms, not only for validated TB drug targets such as DNA gyrase but also for new, yet to be identified, targets.

## INTRODUCTION

Tuberculosis (TB) remains a leading cause of death. Despite global control efforts, the latest World Health Organization (WHO) Global tuberculosis report outlined high incidence rates of multidrug resistant *Mycobacterium tuberculosis* (*Mtb*) strains (MDR-TB), which are resistant to both isoniazid and rifampicin, the two major anti-TB drugs^1^. The control of the current TB-pandemic is hampered by an increase of MDR-TB strains primarily isolated from patients in Eastern Europe and Asia (WHO 2023)^1^. MDR-TB requires treatment with several outdated drugs with important undesirable side effects. Moreover, strains resistant to recently approved drugs are emerging and a high failure rate of compounds in clinical trials requires a constant flow of new lead compounds active against drug-resistant strains.

Major challenges to eradicate TB include discovering and validating new TB targets as well as identifying potent chemical inhibitors with new mechanisms of action and/or new binding sites on validated targets to treat MDR and extensively drug-resistant (XDR) TB. Topoisomerases play crucial roles in several nucleic acid processes and are thus considered as important targets for the development of antimicrobial drugs. In the context of the treatment of TB, current second-line regimens include the use of fluoroquinolones (FQs) (*i.e.* moxifloxacin, gatifloxacin and levofloxacin)^2^, a family of synthetic inhibitors specific to type II bacterial topoisomerases (DNA gyrase and Topo IV). These enzymes are only found in bacteria and share little structural similarities with human topoisomerases, allowing to search for specific inhibitors targeting bacterial topoisomerases without affecting human orthologs^3,4^.

Different from many other bacteria, the mycobacterium genus lacks Topo IV and relies solely on DNA gyrase as a type II topoisomerase, making this enzyme the only known target for FQs^5^. DNA gyrase is the only topoisomerase capable of negative DNA supercoiling, leading to DNA compaction of DNA in bacteria. This enzyme can also catalyse relaxation and (de)catenation in a less efficient way by generating 5’-breaks in double stranded DNA. FQs act as gyrase poisons by blocking the DNA-gyrase complex after binding onto two distinct sites. This type of inhibition can provoke two major cytotoxic effects that ultimately lead to bacterial cell death. On one hand, stabilization of the transient covalent complex formed between the enzyme and the cleaved DNA can generate persistent double-strand breaks, resulting in genome fragmentation. On the other hand, this stabilization prevents the enzyme from completing its catalytic cycle, thereby blocking essential DNA topology-modifying activities such as supercoiling and decatenation. Both mechanisms are believed to contribute to the antibacterial activity of FQs^5,6^. Unfortunately, the continuous emergence of resistant strains to this family of drugs compromises future usage.

One of our long-term objectives is to develop a unified framework to obtain new anti-tuberculous compounds via a primary cell-based screening approach capable of focusing on specific potential targets. The need for the compound to cross the cell membrane to show activity eliminates many false positives that would be obtained from biochemical screens using purified enzyme targets and ensures that the selected agent would be capable of cell penetration, which is required for intracellular targets. We use *Corynebacterium glutamicum* (*Cglu*), a non-pathogenic (BSL-1) surrogate model for *Mtb*, and member of the Corynebacteriales suborder with a short doubling time of under 1h and a genome size comparable to that of *Mtb* (3Mbp). *Cglu* and *Mtb* share core mechanisms related to cell division and cell wall synthesis, as well as a complex multi-layered cell wall^7,8^. Importantly, as written above, *Cglu*, like *Mtb*, encodes only one type II topoisomerase, *i.e.* DNA gyrase.

The active DNA gyrase heterotetramer is constituted of two GyrA and two GyrB subunits (Figure 1)^9,10^. GyrA is composed of an N-terminal region that includes a winged-helix domain (WHD), a tower domain, and a coiled-coil domain, followed by a C-terminal domain (CTD)^11^. In solution, GyrA forms a dimer stabilized by two protein–protein interfaces: one mediated by the WHDs and the other by the globular domains located at the distal ends of the coiled-coil regions^12^. The WHDs harbor the catalytic tyrosine residues responsible for DNA cleavage and religation^12^. The GyrB subunit contains an ATPase domain belonging to the GHKL superfamily (GyrB-Hsp90-histidine/serine protein kinases-MutL), as well as a transducer domain and a topoisomerase-primase (TOPRIM) domain^13,14^. Interaction with GyrA is mediated through contacts between the TOPRIM domains of GyrB and the WHDs of GyrA^15^.

**Figure 1.**
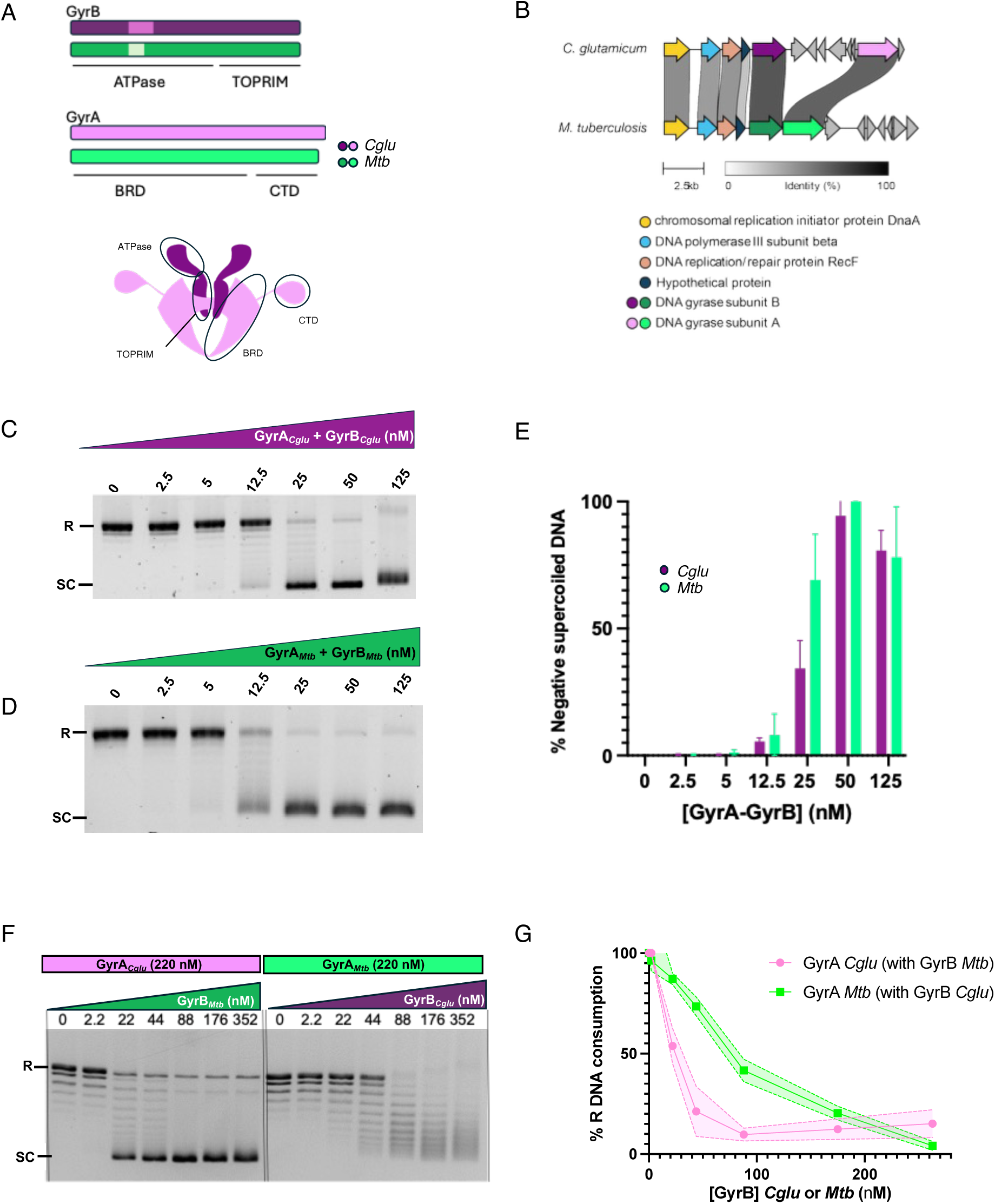
Molecular comparison of DNA gyrase from *Cglu* and *Mtb*: genomes and functional comparison. **A**, Schematic diagram and scheme showing the domain compositions of GyrA and GyrB from *Cglu* (purple) and *Mtb* (green). The GyrA subunit is composed by the ATPase and TOPoisomerase–PRIMase (TOPRIM) domains; the GyrB subunit is composed by the Breakage Reunion Domain (BRD) and the C-terminal Domain (CTD). Within the ATPase domains of GyrB, the species-specific insertion correspondig to the C-loop is depicted in white hues^17,32^. Note that this insertion is absent in other gyrases, such as *E. coli* GyrB (Supplementary figures 4, 5 and supplementary table1). **B**, Clinker-generated^48^ gene cluster showing the positioning of *gyrB* and *gyrA* in *Cglu* and *Mtb* genomes. **C-E**, *In vitro* DNA negative supercoiling activity of *Cglu* and *Mtb* gyrase. Reactions were carried out in the presence of equimolar quantities of GyrA/GyrB subunits on 300 ng relaxed pBR322 plasmid DNA in the presence of 1 mM ATP. Bands corresponding to supercoiled DNA were quantified and reported as percentages for each lane. C-D, Representative activity gels; E, Histograms and error bars represent mean and standard deviations from three independent experiments (n = 3). **F**, Gel of activity assays of negative supercoiled DNA. Supercoiling activity assay on 300 ng as detailed in Methods on relaxed pBR322 plasmid in the presence of 220 nM of GyrA *Cglu* and varying concentrations of GyrB *Mtb* or 220 nM of GyrA *Mtb* in the presence of varying concentrations of GyrB *Cglu*. Negative controls include the presence of sole proteins in the absence of the other subunit. Positive controls included for *Mtb* gyrase (520 nM GyrA+ 197 nM GyrB) and *Cglu* gyrase (85 nM GyrA + 3 μM GyrB) (Supplementary figure 8). **G**, Quantification of relaxed DNA consumption as a function of GyrB concentration. Mean and standard deviation shown for n = 2.

To validate *Cglu* as a model organism for the study of *Mtb* DNA gyrase for drug discovery, we wanted to determine whether the two gyrases are functionally interchangeable. Demonstrating that *Mtb* gyrase can substitute for the native enzyme in *Cglu* would provide strong evidence that *Cglu* constitutes a robust and relevant model for investigating *Mtb* DNA gyrase inhibitors. To address this question, we conducted an integrative analysis of *Cglu* DNA gyrase, both *in vitro* and *in vivo*. Comparative structural, biochemical, and cellular analyses revealed that the two enzymes are highly similar, both catalyzing negative supercoiling, displaying comparable inhibition profiles toward several classes of gyrase-targeting compounds, and sharing conserved drug-binding pockets at the atomic level. At the cellular level, conditional depletion and complementation assays demonstrated that *Cglu* gyrase is essential and that *Mtb* gyrase subunit A (GyrA*_Mtb_*) can restore gyrase function in *Cglu,* highlighting a high degree of functional conservation between the two species. Together, these findings establish *Cglu* as a reliable and non-pathogenic model organism to investigate *Mtb* DNA gyrase function and to accelerate the discovery and improvement of anti-tubercular gyrase inhibitors.

## RESULTS

### Genomic context and sequence features of *Cglu* and *Mtb* DNA gyrases

The active *Cglu* DNA gyrase is a heterotetramer formed by two GyrA and two GyrB subunits encoded by the *gyrA* and *gyrB* genes, respectively (Figure 1A and Supplementary Figures 1,2 and 3). At the genomic level, *Cglu* and *Mtb* differ in their gene organization: in *Cglu*, *gyrB* and *gyrA* are located in distinct operons, separated by six genes (approximately 5 kb), while in *Mtb* the two genes are adjacent within the same operon (Figure 1B and Supplementary Figure 1). Despite this slightly different genomic context, both *Cglu* and *Mtb* are encoded near oriC, where essential genes involved in DNA replication and maintenance are typically clustered^16^. Nevertheless, the encoded GyrB and GyrA subunits are highly similar between the two species, highlighting the strong evolutionary conservation of this essential enzyme (Figure 1 and Supplementary Table 1). They share about 70% sequence identity and all critical residues involved in substrate binding and catalysis are strictly conserved (Supplementary Figures 2, 3 and Supplementary Table 1)^11^. The lower sequence conservation is mainly localized at the GyrB N-terminus and the GyrA C-terminus region, two regions that are longer in *Cglu* than in *Mtb* but retain similar physico-chemical properties (Supplementary Table 1 and Supplementary Figure 4)^17,18^. Also, both enzymes contain the Corynebacteriales*-*specific C-loop within the ATPase domain, located between the last two β-strands of the GHKL domain^17^. Although this insertion exhibits variable amino acid sequences among Corynebacteriales, it preserves a characteristic acidic composition (Supplementary Table 1)^17^. Associated to this C-loop, the DEEE-loop in the GyrA-BRD domain, known to be crucial in *Mtb* gyrase to stabilize the resting enzyme through interactions with the C-loop^17^, corresponds in *Cglu* to an EESE-loop (residues 211-214 in *Mtb* / 208-211 in *Cglu*), and is likely to exert a similar structural role (Supplementary Figure 4).

### Functional features of *Cglu* and *Mtb* DNA gyrases

We produced N-terminally tagged recombinant GyrA*_Cglu_* and GyrB*_Cglu_* subunits for biochemical and structural characterization (Supplementary Figure 5A and Supplementary Table 2). After overexpression in *E. coli*, the proteins were purified using a three-step purification procedure including removal of the His-SUMO tag. Size-exclusion chromatography (SEC) showed that GyrA*_Cglu_* and GyrB*_Cglu_* each eluted as a single peak (Supplementary Figure 5B, C). For comparison, *Mtb* gyrase was purified following the protocol that we previously established and validated^19^, ensuring consistency in purification and activity assessment between the two enzymes. In most prokaryotes, the two essential DNA topological activities, negative DNA supercoiling and decatenation, are carried out by distinct type IIA topoisomerases, DNA gyrase and topoisomerase IV, respectively^8^. In *Mtb*, however, gyrase is the sole type IIA topoisomerase and catalyzes both functions *in vivo* ^20,21^. To compare enzymatic activities, we first defined reaction conditions supporting DNA supercoiling activity for both gyrases. We found that the buffer previously optimized for monitoring *Mtb* gyrase activity^21^ was also suitable for *Cglu* gyrase, thus, all subsequent experiments were performed using these conditions at 37°C. Both enzymes were found to catalyze full negative supercoiling of 50 nM of relaxed pBR322 plasmid (Figure 1C-E). Furthermore, decatenation assays validated that both *Cglu* and *Mtb* gyrases are capable of decatenating kinetoplast DNA substrate. For *Cglu* gyrase a detectable activity was observed at approximately 50 nM, whereas *Mtb* gyrase required a five-fold higher concentration for comparable activity (Supplementary Figure 6).

Next, we asked whether heterologous combinations of the two subunits (GyrB, GyrA) could be functional. Within the gyrase heterotetramer, the principal contacts between the subunits occur between the GyrB-TOPRIM and the GyrA-BRD domains (Figure 1A). We therefore used AlphaFold3 (AF3) to generate the heterologous models 2GyrB-TOPRIM*_Cglu_* / 2GyrA-BRD*_Mtb_* and 2GyrB-TOPRIM*_Mtb_* / 2GyrA-BRD*_Cglu_*, and compared them with the respective WT forms (Supplementary Figure 7). Interface analysis with the PISA server identified largely conserved contact regions, with only minor residue variations in the heterologous combinations (Extended data Tables 1 to 4). The predicted confidence scores for all models were very high (>0.9) and comparable to those obtained for the WT enzyme, suggesting that the heterologous gyrases remain functional. The GyrB-TOPRIM and the GyrA-BRD interaction pattern is virtually identical in *Cglu* and *Mtb*, with all major interface residues participating in equivalent contacts despite the minor differences (Supplementary Figure 7I). Accordingly, the purified proteins GyrA*_Mtb_* / GyrB*_Cglu_* and GyrA*_Cglu_* / GyrB_Mtb_ were both shown to catalyze DNA supercoiling, albeit with distinct efficiencies (Figures 1F-G, Supplementary Figure 8). As a whole, while AF3 predictions indicated robust and conserved interaction interfaces between *Mtb* and *Cglu* subunits, experimental data uncovered some functional asymmetry between the two heterologous combinations, with the GyrA*_Mtb_*/GyrB*_Cglu_* complex showing reduced catalytic activity possibly due to inherent differences in catalytic properties.

### Conditional depletion of GyrA or GyrB induces strong morphological changes in *Cglu*

As the DNA gyrase is essential for bacterial survival, we constructed two conditional mutant strains where the transcription of either *gyrA* or *gyrB* was uncoupled from its native promoter. A transcriptional terminator was placed before the *gyrA* or *gyrB* genes, and the transcription of each gene was put under the control of the previously described *myo-*inositol repressible promoter (*P_ino_*) of the inositol phosphate synthase *ino1* gene (Figure 2A)^22^ to yield the *P_ino_-gyrA* and *P_ino_-gyrB* mutant strains. No downstream polar effects were expected, as *gyrA* and *gyrB* reside in distinct operons: *gyrB* is the terminal gene in its operon, and *Cgl0014* (the first gene after *gyrA*) possesses its own promoter (Supplementary Figure 1). Depletion of GyrA*_Cglu_* or GyrB*_Cglu_* was confirmed by Western blot, where the corresponding protein signal disappeared after growth in medium containing 1% *myo-*inositol (Figure 2B and Supplementary Figure 9). As anticipated, both depleted strains were non-viable on solid medium supplemented with 1% *myo-*inositol (Figure 2C). To investigate the cellular consequences of gyrase depletion, we examined the morphology of the *P_ino_-gyrA* and *P_ino_-gyrB* strains under repressing conditions (1% *myo-*inositol). A first noticeable feature was the emergence of highly heterogeneous cell population, similar for both strains, with the most striking effect observed on nucleoid morphology (Figures 2D-E). Heatmaps of DNA foci distribution revealed three distinct cell populations compared with the WT: anucleate cells lacking detectable DNA, cells with a single loose nucleoid, and cells displaying one to three more compacted nucleoids (Figure 2E). Nucleoid decompaction can lead to bulged poles (Figure 2D). The gyrase-depleted cells were longer and wider than the WT, with the most notable effect on cell width (Figures 2F-G). Scatter plots of cell length versus DNA foci number confirmed the presence of anucleate cells in both depleted strains with a proportion of 3% for *P_ino_-gyrA* and 6% for *P_ino_-gyrB*, indicating a partial defect in DNA replication or segregation (Supplementary Figure 10).

**Figure 2.**
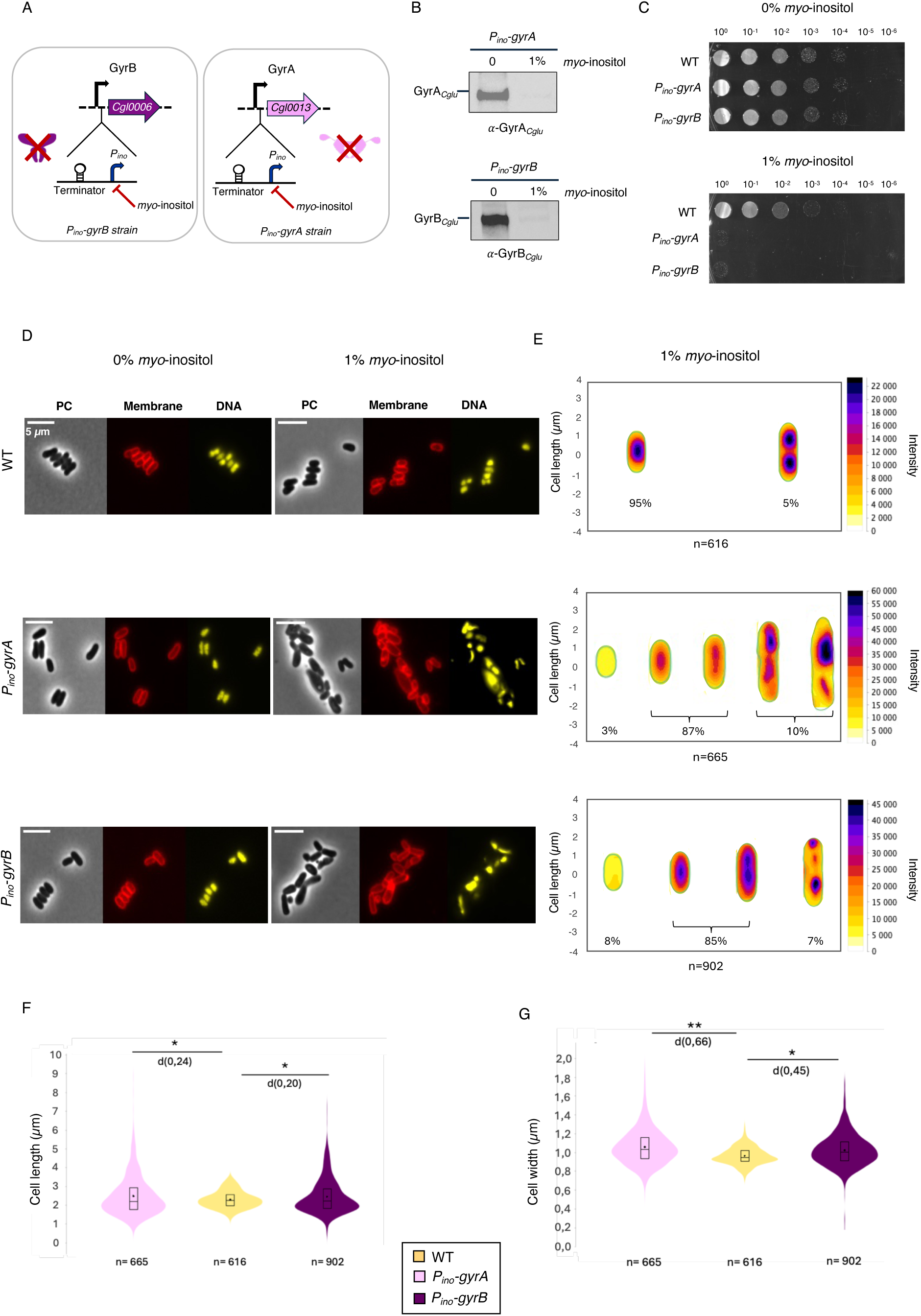
Construction of conditional *P_ino_-gyrA* and *P_ino_-gyrB Cglu* strains and phenotypic characterization of silencing effect. **A**, Principle of *P_ino_* conditional knock-out system in *Cglu*. Specific genomic insertions of a *P_ino_* promoter before *gyrB* or *gyrA* ORFs allowed the generation of *P_ino_*-*gyrB* or *P_ino_*-*gyrA Cglu* strains respectively. Selective silencing of translation of GyrB or GyrA was therefore possible by addition of 1% *myo*-inositol in culture medium. **B**, Spot assay for growth phenotypes of wild-type ATCC1332 (WT), *P_ino_*-*gyrB* and *P_ino_-gyrA Cglu* strains on CGIIX medium agar in the absence or presence of 1% *myo*-inositol. Overnight cultures of each *Cglu* strain in CGIIX medium with or without 1% *myo*-inositol were normalized to an OD_600_ of 0.5 and deposited as serially diluted 10-fold spots. The number of cells in each spot is indicated above. **C**, Western blots of whole cell extracts from the *P_ino_-gyrA* and *P_ino_-gyrB* strains, in the absence or presence of 1% *myo*-inositol. GyrA and GyrB levels were revealed using anti-GyrA*_Cglu_* (α-GyrA*_Cglu_*) or anti-GyrB*_Cglu_* (α-GyrB*_Cglu_*) antibodies. Molecular weights (MW) are indicated in the uncropped WB (Supplementary Figure 9 + Extended data Figure 1). **D**-**E**, (**D**), Representative images in phase contrast (PC), membrane staining (Nile Red) and DNA (Hoechst) staining signals at 24 hours of cells grown in controlled medium CGXII supplemented with 4% sucrose in the absence or presence of 1% *myo-*inositol. Scale bars correspond to 5 μm. (**E**), Corresponding heatmaps representing the localization pattern of DNA after 24 hours. n numbers represent the number of cells used in the analyses. Percentages of bacteria exhibiting each DNA localization are shown below each heatmap.The data shown are representative of experiments made independently in triplicate. **F**, Violin plots illustrating the distribution of cell length for each tested condition from panel D at 24 hours after *myo-*inositol addition for cells grown in controlled medium CGXII supplemented with 4% sucrose. (Cohen’s d, from left to right compared to control: (*, d(*P*_ino-_ *gyrA)* = 0.24, d(*P*_ino-_ *gyrB*) = 0.20). The box indicates the 25th to 75th percentiles, and whiskers indicate the 95% confidence interval. The mean and median are represented by a dot and a line within the box. The number of cells (n) taken into account for the analysis is indicated below each violin. **G**, Violin plots illustrating the distribution of cell width for each tested condition from panel D at 24 hours after *myo*-inositol addition for cells grown in controlled medium CGXII supplemented with 4% sucrose (Cohen’s d, from left to right compared to control: (**,d(*P_ino-_gyrA)* = 0.66; *, d(*P_ino-_gyrB*) = 0.45). The box indicates the 25th to 75th percentiles, and whiskers indicate the 95% confidence interval. The mean and median are represented by a dot and a line within the box. The number of cells (n) taken into account for the analysis is indicated below each violin.

Because depletion of GyrA*_Cglu_* or GyrB*_Cglu_* produced nearly identical phenotypes, in both cases linked to the absence of a functional DNA gyrase heterotetramer, we focused subsequent analyses on the *P_ino_-gyrA* strain, as most FQ resistance mutations in *Mtb* are located in the GyrA subunit^23^. To verify that observed phenotypes were solely due to GyrA*_Cglu_* depletion and not to potential polar effects on neighboring genes, we performed a complementation assay by expressing GyrA*_Cglu_* from a pTGR vector under the control of the stringent gluconate promoter *P_gntK_*, which is repressed by 4% sucrose and induced by 1% gluconate^24,25,26^. Unless otherwise specified, subsequent experiments used minimal medium (CGXII, see Materials and Methods) supplemented with either 4% sucrose; 4% sucrose + 1% *myo*-inositol; or 4% sucrose + 1% *myo*-inositol + 1% gluconate. The empty plasmid introduced in the *P_ino_-gyrA* strain (*P_ino_-gyrA + P_gntK_-empty*) was used as control. The growth of the complemented *P_ino_-gyrA* strain was evaluated by serial-dilution spot assays on CGXII plates in the presence of 50 μg/ml kanamycin.

Complementation experiments were done by expressing a fluorescent version of GyrA*_Cglu_* (*P_gntk_*-*gyrA-mNeon*) in the *P_ino_-gyrA* strain (Figure 3A). Western blot analysis confirmed the full depletion of the endogenous GyrA*_Cglu_* in the presence of 1% *myo*-inositol, while GyrB*_Cglu_* was detected in both control and complemented strains (Figure 3B and Supplementary Figure 11). The presence of GyrB*_Cglu_* under GyrA*_Cglu_* depletion conditions indicates that the two subunits are expressed independently and stably, and that complementation with GyrA alone should be sufficient to restore a normal phenotype. The functional restoration of gyrase activity was demonstrated by spot dilution assays, as even in sucrose conditions (i.e. minimal promoter leakage) the complemented strain recovered growth under repression conditions (1% *myo*-inositol, 4% sucrose), while the strain carrying the empty vector was unable to grow (Figure 3C). Moreover, both the control and the complemented *P_ino_-gyrA +* GyrA*_Cglu_*-mNeon strains exhibited complete recovery of the WT morphology, as indicated by heatmaps of DNA foci, and violin plots of cell length and cell width (Figure 3D-F) and DNA content (Supplementary Figure 10). Interestingly, the ectopic mNeon fusion localized in the cytoplasm as expected, distributed along the bacterial chromosome (Figure 3D). Taken together, these results demonstrate that the phenotypes observed in the *P_ino_-gyrA* strain are directly attributable to GyrA*_Cglu_* depletion and not to off-target or polar effects.

**Figure 3.**
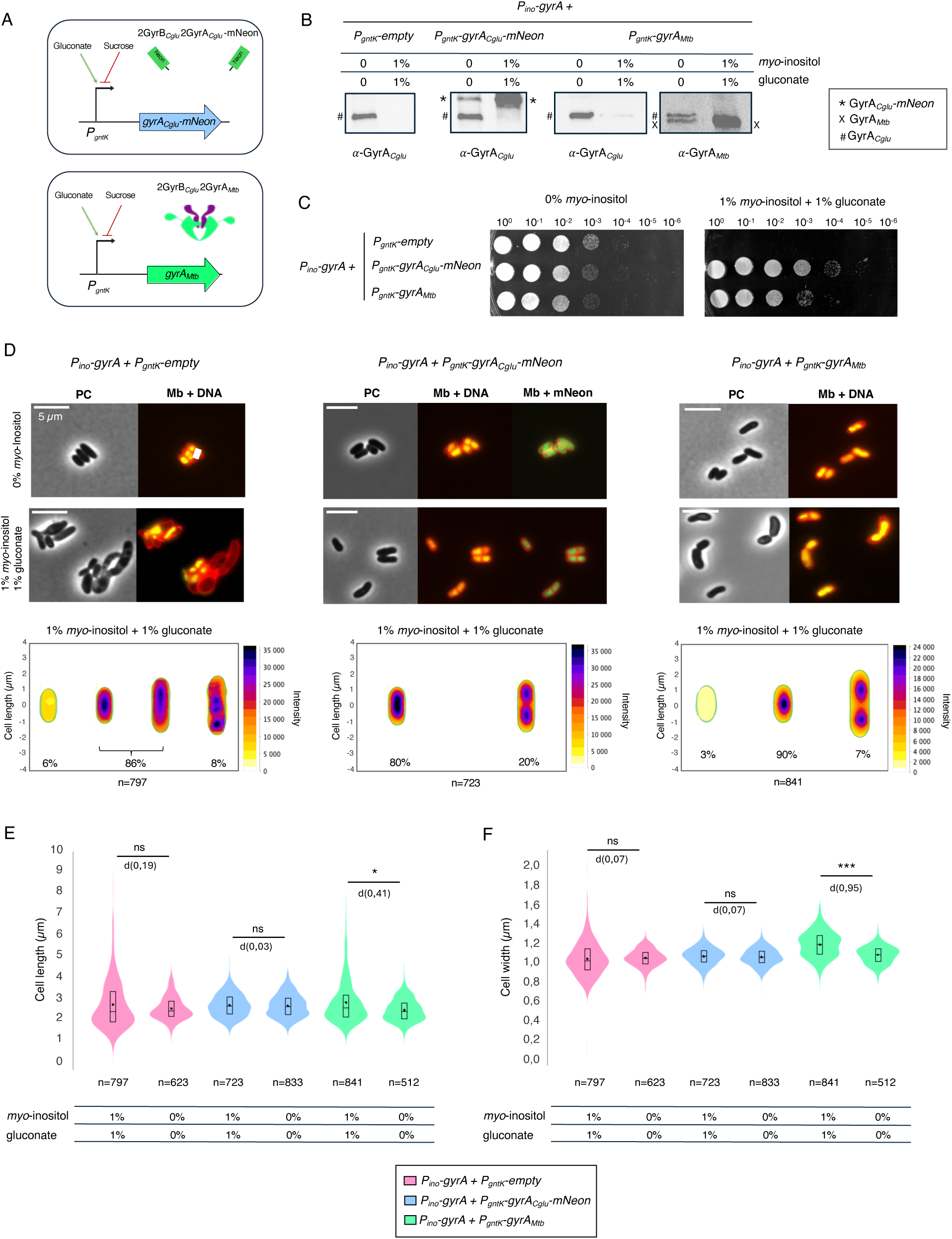
Phenotypic characterization of *P_ino_-gyrA* complemented with GyrA*_Cglu_*-mNeon or GyrA*_Mtb_*. **A**, Principle of complementation of *P_ino_-gyrA* strain with GyrA*_Cglu_*-mNeon or GyrA*_Mtb_*. *P_ino_-gyrA strain* was transformed with plasmid pTGR harboring *gyrA_Cglu_*-mNeon or *gyrA_Mtb_* under the control of the promoter *P_gntk_*. The latter element enables selective activation or inhibition by addition of gluconate or sucrose in culture medium. **B**, Western blots of whole cell extracts from the *P_ino_-gyrA* + *P_gntk_-empty* plasmid or + *P_gntk_-gyrACglu-mNeon* plasmid or + *P_gntk_- gyrA_Mtb_* plasmids strains, in the absence or presence of 1% *myo*-inositol + 1% gluconate. GyrA*_Cglu_*, GyrA*_Cglu_*-mNeon and GyrA*_Mtb_* levels were revealed using anti-GyrA*_Cglu_* (α-GyrA*_Cglu_*), anti-mNeon (α-GyrB*_Cglu_*) or anti-GyrA*_Mtb_* (α-GyrA*_Mtb_*) antibodies. Molecular weights (MW) are indicated in the uncropped WB (Supplementary Figure 11 + Extended data Figure1). **C**, Spot assay for growth phenotypes of *P_ino_-gyrA Cglu* complemented with empty vector, with g*yrA_Cglu_-mNeon* or with g*yrA_Mtb_* carrying vector on controlled medium agar in the absence or presence of 1% *myo*-inositol and 1% gluconate. Overnight cultures of each *Cglu* strain in controlled medium with or without 1% *myo*-inositol and 1% gluconate were normalized to an OD_600_ of 0.5 and deposited as serially diluted 10-fold spots. **D**, Representative images in phase contrast (PC), membrane staining (Nile Red), DNA (Hoechst) and mNeon green fluorescence at 24 hours after *myo-*inositol and gluconate addition for cells grown in controlled medium CGXII supplemented with 4% sucrose. Heatmaps represent the localization pattern of DNA after 24 hours. n numbers represent the number of cells used in the analyses. Scale bars correspond to 5 μm. Percentages of bacteria exhibiting each DNA localization are shown below each heatmap.The data shown are representative of experiments made independently in triplicate. **E-F,** Violin plots illustrate cell length and cell width distributions for the depletion strain *P_ino-_gyrA + P_gntK_-empty,* the complementation strain *P_ino-_gyrA + P_gntK_ gyrA-mNeon* and the complementation strain *P_ino-_gyrA + P_gntK_-gyrA_Mtb_* grown at 24 hours with or without *myo*-inositol and gluconate for cells grown in controlled medium CGXII supplemented with 4% sucrose. **E,** Violin plots illustrating the distribution of cell length for each tested condition from panel D at 24 hours with and without *myo-*inositol and gluconate addition for cells grown in controlled medium CGXII supplemented with 4% sucrose. Cohen’s d, between *P_ino-_gyrA + P_gntK_-empty* with and without *myo*-inositol and gluconate: (ns,d= 0.19), Cohen’s d, between *P_ino-_gyrA + P_gntK_-gyrA-mNeon* with and without *m*yo-inositol and gluconate: (ns,d= 0.03), Cohen’s d, between *P_ino-_gyrA + P_gntK_-gyrA_Mtb_* with and without myo-inositol and gluconate: (*, d= 0.41). **F,** Violin plots illustrating the distribution of cell width for each tested condition from panel D at 24 hours with and without *myo-*inositol and gluconate addition for cell grown in controlled medium CGXII supplemented with 4% sucrose. Cohen’s d, between *P_ino-_gyrA + P_gntK_-empty* with and without *myo*-inositol and gluconate: (ns, d= 0.07), Cohen’s d, between *P_ino-_gyrA + P_gntK_-gyrA-mNeon* with and without *myo*-inositol and gluconate: (ns, d= 0.07), Cohen’s d, between *P_ino-_gyrA + P_gntK_-gyrA_Mtb_* with and without *myo*-inositol and gluconate: (***, d= 0.95). The box indicates the 25th to 75th percentiles, and whiskers indicate the 95% confidence interval. The mean and median are represented by a dot and a line within the box. The number of cells (n) taken into account for the analysis is indicated below each violin.

### Interchangeability of GyrA subunits from *Cglu* and *Mtb*

To investigate whether GyrA*_Mtb_* could functionally replace its GyrA*_Cglu_*, we performed the same experiments as described above, this time complementing the *P_ino_-gyrA* depleted strain with a *P_gntk_*-*gyrA_Mtb_* plasmid (Figure 3A). The depletion of the endogenous GyrA*_Cglu_* subunit was confirmed by Western blot analysis in the complemented strain, and the induced expression of GyrA*_Mtb_* and native levels of GyrB*_Cglu_* were assessed (Figure 3B and Supplementary Figure 12). The expression of GyrA*_Mtb_* successfully restored normal growth on solid media (Figure 3C). Cells exhibited a mostly normal DNA organization compared to the depleted strain (Figure 3D), typically displaying one or two distinct and well-compacted DNA foci, similar to the WT (Figures 2E and 3D).

Although the complemented cells remained slightly elongated on average compared to WT cells (Figure 3E), they exhibited full restoration of viability and proper DNA segregation, as reflected by their nucleoid organization and the disappearance of most anucleate cells (Figure 3D and Supplementary Figure 13). In line with the partial structural compatibility of the heterologous enzyme, the GyrA*_Mtb_*/GyrB*_Cglu_* complex displayed reduced activity *in vitro*. Nevertheless, the successful rescue of growth and overall cell organization demonstrates that *Mtb* gyrase can efficiently support essential processes in *Cglu*.

Having established that the gyrase activities are at least partially interchangeable between *Mtb* and *Cglu*, we next sought to examine more closely how different classes of anti-gyrase inhibitors influence *Cglu* gyrase activity and cell morphology, and whether this functional compatibility extends to the conservation of drug-binding sites.

### Phenotypic effects of anti-gyrase compounds

To assess the *in vitro* susceptibility of *Cglu* DNA gyrase to known inhibitors, we determined the IC_50_ values for negative supercoiling inhibition of three different anti-gyrase compound classes (Table 1). The lowest IC_50_ value was obtained for novobiocin, an aminocoumarin that specifically targets the ATPase domain of gyrase. Moxifloxacin (MFX) and ciprofloxacin (CIP), the two fluoroquinolones that we evaluated, exhibited comparable inhibitory potencies, with the IC_50_ for MFX being approximately twofold lower than that of CIP (Table 1). Interestingly, gepotidacin, a member of the Novel Bacterial Topoisomerase Inhibitor (NBTI) class^27^, showed an IC_50_ similar to that of MFX (Table 1). Comparison with previously reported IC_50_ values for *Mtb* DNA gyrase revealed broadly comparable susceptibility profiles across the two enzymes (Table 1). In particular, both gyrases were more sensitive to novobiocin and displayed similar relative sensitivities to FQ and non-FQ inhibitors, consistent with previously reported MIC data showing a similar overall trend in the two species^28^. Given their strong biochemical resemblance, *Cglu* would therefore provide a useful model to explore the cellular consequences of gyrase inhibition and to discriminate the morphological effects induced by different inhibitor classes. To investigate how representative inhibitors from each class affected *Cglu* cell and nucleoid morphology, cells were exposed to each compound at 10X MIC in order to maximize phenotypic differences. In terms of cell width, FQ-treated cells remained similar to untreated WT cells, whereas novobiocin produced a larger width distribution (0.3-1.7 µm; mean 1.1-1.3 µm) and more heterogeneous morphologies, including localized swelling and bulging (Figure 4 and Supplementary Figure 14). All tested compounds promoted cell elongation, which was particularly pronounced following treatment with the FQs ciprofloxacin and moxifloxacin, and to a lesser extent with novobiocin (Figure 4). Scatter plot analyses of cell length versus number of nucleoids further highlighted differences between compound classes (Supplementary Figure 15). Markedly elongated CIP- and MFX-treated cells often contained 0-3 DNA foci, indicative of defects in chromosome replication and segregation. In contrast, novobiocin-treated cells also exhibited elongation but displayed fewer anucleate cells, consistent with a predominant replication defect rather than complete segregation failure. To further confirm the overall similarity in inhibition profiles between *Cglu* and *Mtb* gyrases, we next explored the structural basis of inhibitor binding at atomic resolution.

**Figure 4.**
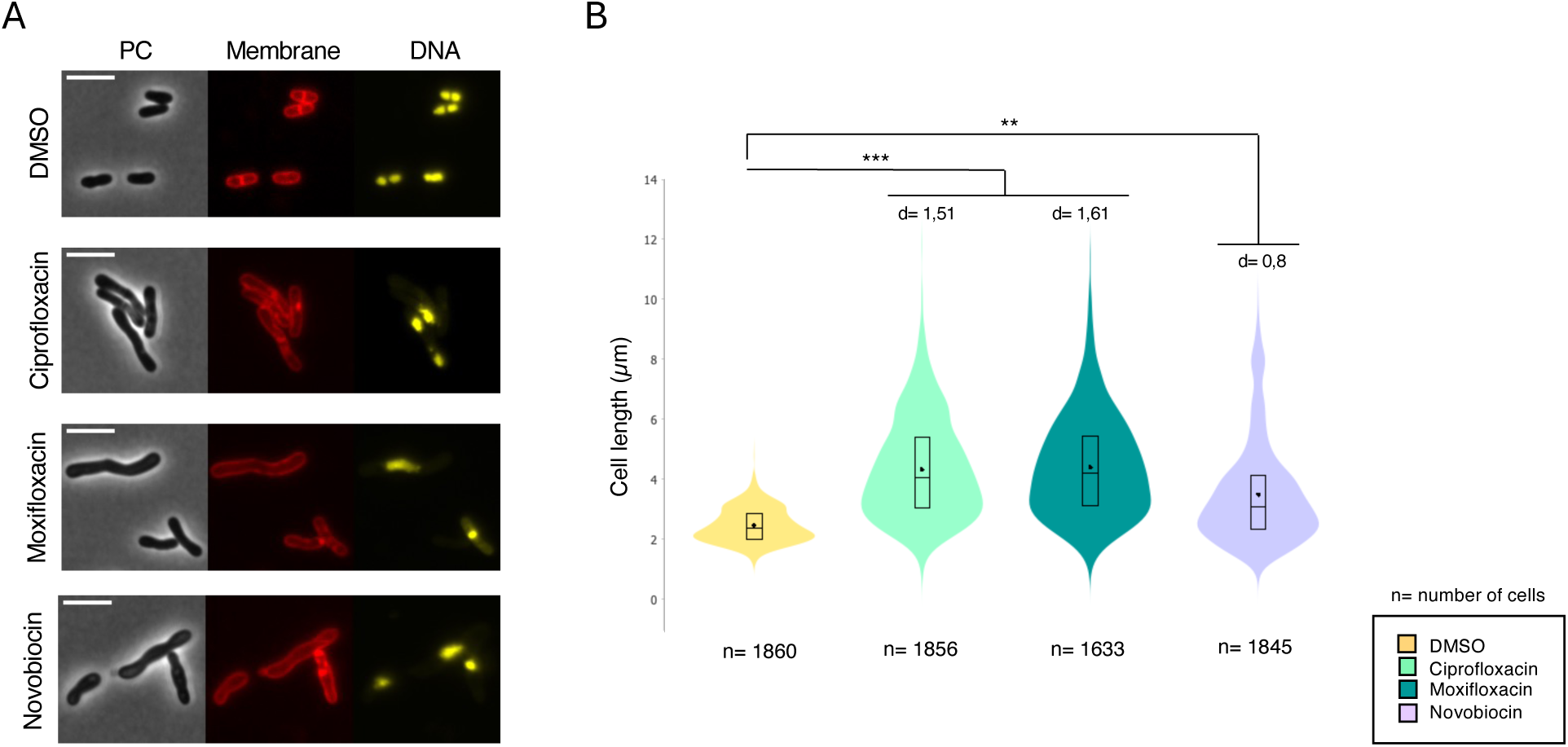
Evaluation of the impact of Ciprofloxacin, Moxifloxacin and Novobiocin on the morphotype of *Cglu* ATCC13032. **A**, Representative images in phase contrast (PC), membrane staining (Nile Red) and DNA (Hoechst) of *Cglu* WT in the absence (DMSO, control) or in the presence of each indicated drug. Scale bars correspond to 5 μm. **B**, Violin plots illustrating the distribution of cell length for each tested condition from **A**. Cohen’s d, from left to right compared to control: (***, d(Ciprofloxacin) = 1.51, d(Moxifloxacin) = 1.61; **, d(Novobiocin) = 0,8). The box indicates the 25th to 75th percentiles, and whiskers indicate the 95% confidence interval. The mean and median are represented by a dot and a line within the box. The number of cells (n) taken into account for the analysis is indicated below each violin.

**Table 1.** IC_50_ values of antibiotics targeting DNA gyrase.

| **Antibiotic \ Species** | **IC<sub>50</sub> (μM) *Cglu*** | **IC<sub>50</sub> (μM) *Mtb*** |
| --- | --- | --- |
| **Moxifloxacin** | **8.9 (+/- 5.4)** | **5<sup>49</sup>** |
| **Ciprofloxacin** | **16.6 (+/- 7.2)** | **25<sup>19</sup>** |
| **Novobiocin** | **0.52 (+/- 0.23)** | **0.49<sup>50</sup>** |
| **Gepotidacin** | **5.4 (+/- 0.6)** | **15<sup>19</sup>** |

### The inhibitor binding pockets are highly conserved between *Cglu* and *Mtb* gyrases

The dynamic nature of the gyrase, which undergoes large conformational changes to accommodate DNA binding and cleavage, makes structural analysis challenging. As previously described for the *Mtb* gyrase to prevent subunit dissociation and generate a more stable biological unit^17^, we linked GyrB*_Cglu_* and GyrA*_Cglu_* into a single polypeptide chain (termed GyrBA*_Cglu_*) leading to a functional dimeric enzyme (GyrBA*_Cglu_*; ∼170 kDa per monomer; Supplementary Figure 5 and Supplementary Table 1). As expected, the fused gyrase remained active and displayed approximately twice the supercoiling activity of the non-fused subunits (Supplementary Figure 16). Cryo-EM analysis of GyrBA*_Cglu_*_2_ bound to 150 bp dsDNA and moxifloxacin yielded a density map of the cleavage core (two BRD-TOPRIM dimers) at an overall resolution of 3.2 Å (Figure 5A, supplementary Figure 17 and Supplementary Table 3). CTDs and ATPase domains were visible at lower resolution (>5 Å) and not modelled (Figure 5A). Like other bacterial type IIA topoisomerase–DNA complexes, the *Cglu* cleavage core binds a single dsDNA bridging the dimer interface (Figures 5A)^29,30,31^. Each strand is cleaved once, with the 5ʹ-phosphate group covalently linked to the catalytic Tyr126 of GyrA*_Cg_*, equivalent to Tyr129 in *Mtb* (Supplementary Figure 18). The presence of the magnesium ions coordinated by Glu495, Asp568, and Asp570 of GyrB close to the Tyr_cat_ was also identified (equivalent to Asp532, Asp534, Glu459 in *Mtb,* Supplementary Figure 18)^30^. Furthermore, density maps clearly revealed two moxifloxacin molecules positioned between the cleaved DNA ends on both halves of the complex (Supplementary Figures 18 and 19). Each molecule binds within a pocket primarily shaped by DNA base stacking and is further stabilized via a magnesium ion coordinated by the keto-acid group of each quinolone core (Figure 5B and Supplementary Figure 18). Superposition of the *Cglu* gyrase structure and the crystallographic *Mtb* cleavage core in complex with moxifloxacin shows structural conservation (RMSD of 1.4 Å for all Cα atoms), confirming that FQs trap both enzymes in the same closed conformation (Supplementary Figure 19).

**Figure 5.**
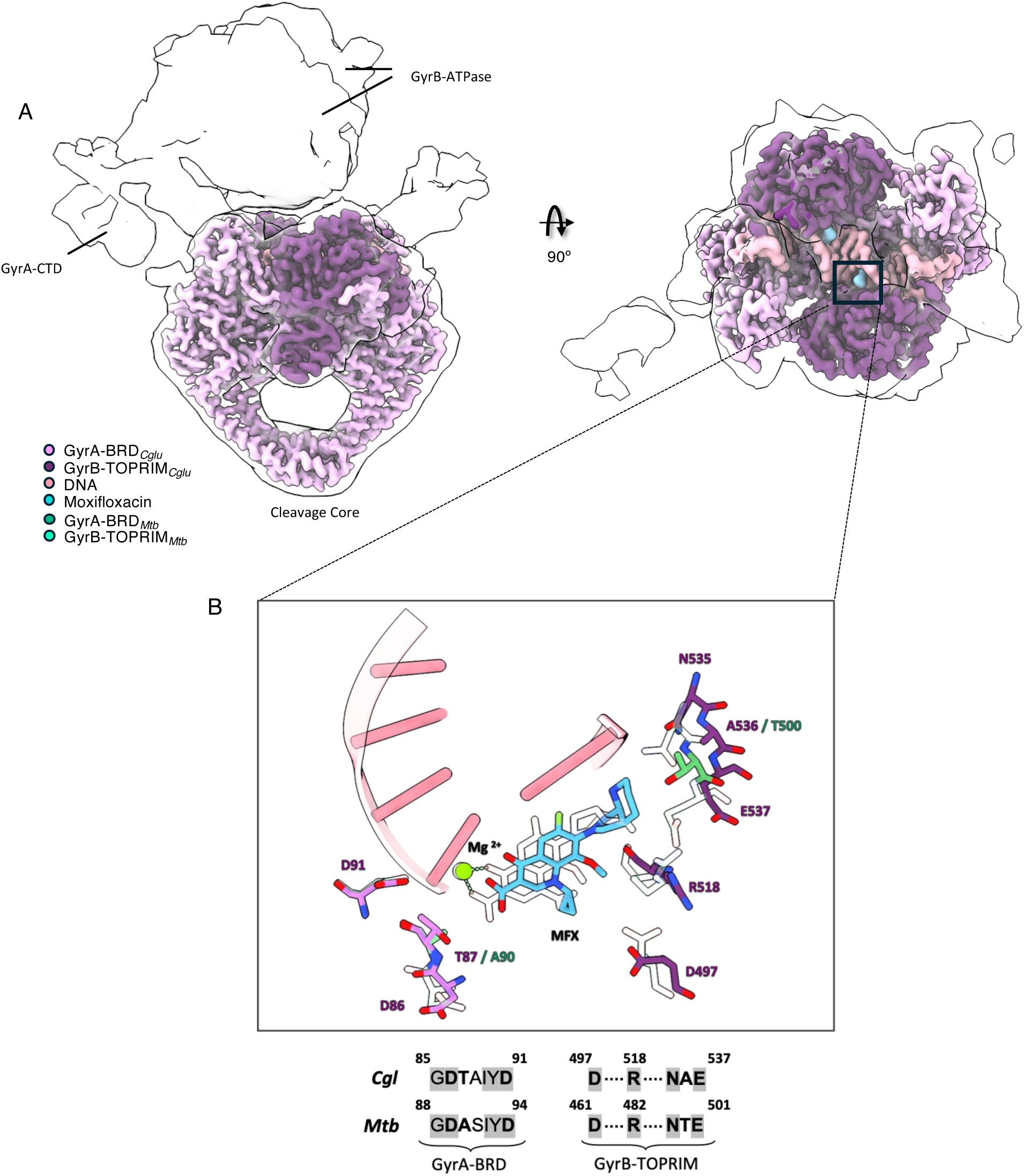
Molecular comparison of DNA gyrase from *Cglu* and *Mtb*: 3D structures comparison and analysis of the moxifloxacin binding pocket. **A**, Cryo-EM maps of *Cglu* gyrase in complex with dsDNA and two molecules of moxifloxacin (MFX). A low-resolution map (5 Å) illustrates the position of the GyrA CTDs and GyrB ATPase domains. The high-resolution (3.2 Å) cleavage core map (two GyrB-TOPRIM and two GyrA-BRD) is coloured according to scheme in Figure 1A: GyrA-BRD in light purple and GyrB-TOPRIM in dark purple. Ligands DNA and MFX are shown in pink and cyan. Low-resolution cryo-EM map is in white with 60% of transparency. **B,** Zoom on MFX binding pockets in *Cglu* and *Mtb* gyrase from superposition shown in Supplementary Figure 19A. Mg^2+^ ions are in light green sphere representation. *Mtb* gyrase is coloured in green.

The detailed comparison of the two MFX-binding pockets indicates that all but two residues close to the bound ligand are strictly conserved between the *Cglu* and *Mtb* gyrases. Divergent positions are Thr87 (Ala90 in *Mtb*) near the Mg^2+^ important for FQs stabilization, and Ala538 (Thr500 in *Mtb*) in GyrB*_Cglu_* (Figure 5B). Residue 90 in *Mtb* is a well-known determinant of FQs sensitivity: when a serine occupies this position, like in *E. coli* gyrase, the drug is better stabilized, whereas an alanine in this position, as found in *Mtb*, reduces affinity^30,32^. In *Cglu*, the corresponding residue is a threonine (Thr87), positioned close to the Mg^2+^ ion essential for FQs stabilization, that likely provides an intermediate level of stabilization and slightly shifts moxifloxacin within the pocket. On the opposite side, the replacement of *Mtb* Thr500 by *Cglu* Ala538 induces a movement of the adjacent loop facilitating antibiotic accommodation (Figure 5B). Together, these two compensatory substitutions appear to maintain optimal moxifloxacin binding in *Cglu* gyrase despite minor local rearrangements of the binding pocket (Figure 5B).

Although no structures were determined experimentally for *Cglu* gyrase bound to other inhibitors, conservation analyses of the corresponding binding regions revealed that key residues forming the novobiocin and NBTI binding pockets are also conserved between *Cglu* and *Mtb*. Superposition of the *Cglu* cryo-EM structure with that of *Mtb* bound to BDM71403 (an NBTI analog to gepotidacin)^19^ showed that the inhibitor occupies a pocket framed by DNA base pairs on the two-fold axis and residues from GyrA-BRD domains (Supplementary Figure 19B). Similarly, the novobiocin-cavity, located in the ATPase (GHKL) domain, overlaps with the ATP site and includes residues involved in Mg^2+^ coordination and hydrogen-bonding with ATP phosphates and adenine. Structural modeling based on *Thermus thermophilus* gyrase complexed with novobiocin^33^ confirmed that the composition of this pocket is strictly conserved in *Mtb* and *Cglu* (Supplementary Figure 19C).

Together, the above analyses indicate that *Cglu* gyrase shares with *Mtb* not only an overall structural architecture but also highly conserved binding pockets for antibiotics targeting both the cleavage and ATPase cores, supporting the relevance as a robust surrogate for mechanistic and inhibitor studies.

## DISCUSSION

In this work, we validated *Cglu* as a non-pathogenic surrogate for *Mtb* to facilitate the discovery of novel DNA gyrase inhibitors through comparative structural and functional analyses. To functionally dissect gyrase in this model, we generated cKD mutant strains for the *Cglu* gyrase subunits, GyrA*_Cglu_* and GyrB*_Cglu_*. As expected for an enzyme that maintains DNA topology, depletion of either subunit resulted in severe growth impairment and characteristic morphological abnormalities, confirming that gyrase activity is indispensable for *Cglu* viability. Our work provides description of the morphotypes associated with gyrase depletion in this species and within the Corynebacteriales. Previous studies on *Mtb* and *Mycobacterium smegmatis* (*Msm*) described the essential nature of gyrase and reported comparable cell-elongation and nucleoid-organization defects upon inhibition or depletion of the enzyme^34,35^, and the present work complements these findings with a more detailed spatial and quantitative morphometric characterization. Through imaging and quantitative morphometric analyses, we showed that both *P_ino_*-*gyrA* and *P_ino_*-*gyrB* strains displayed similar phenotypic outcomes including distinct subpopulations within repressed cultures, anucleate cells, cells with decompacted DNA and cells with one or several DNA foci, revealing a heterogeneity in the cellular response to gyrase loss (Figure 2). Beyond these morphological alterations, the observed defects reflect the central role of the gyrase in maintaining global DNA topology, which directly influences the proper functioning of the replication, transcription and segregation machineries. Perturbations in supercoiling homeostasis are expected to generate conflicts between replication and transcription, cause nucleoid disorganization and disrupt cell division, all consistent with the phenotypes observed here. Crucially, complementation of the *P_ino_*-*gyrA* strain with a functional, fluorescently tagged GyrA-mNeon variant not only restored normal growth, nucleoid structure and cell morphology, but also enabled direct visualization of gyrase distribution within the cytoplasm (Figure 3). The homogeneous fluorescence signal throughout the nucleoid region confirms that the gyrase is evenly distributed along the bacterial chromosome, consistent with its dynamic action on supercoiling throughout the genome. Together, these results position *Cglu* as a robust model to dissect gyrase-dependent control of chromosome architecture and to visualize, at unprecedented resolution, the cellular consequences of disrupting DNA topology in Actinobacteria.

Building on these genetic observations, we next compared gyrase depletion with chemical inhibition to explore how different perturbations modes translate into distinct morphological outcomes. The comparative analysis between this work and previous studies^19^ highlights both shared and mechanistically distinct morphological signatures. Across all gyrase inhibitors tested, FQs (ciprofloxacin, moxifloxacin), the aminocoumarin novobiocin and the NBTI-class compounds, cells displayed elongation and visible perturbations of nucleoid architecture, consistent with the central role of gyrase in maintaining DNA topology. However, the extent and nature of these alterations varied markedly depending on the inhibitor class, reflecting different modes of enzyme engagement and downstream cellular responses. Among all conditions, FQs treatment produced the most extreme phenotype, with pronounced filamentation and severe nucleoid condensation or loss of detectable DNA foci. A significant fraction of ciprofloxacin-treated cells exceeded 8 μm in length, far greater than those observed in either novobiocin- or NBTI-treated cultures. While this morphology overlaps with that of the GyrA- and GyrB-depleted strains, the increased severity of the FQ-induced phenotype likely reflects double-strand DNA breaks (DSBs). These breaks occur when the antibiotic traps gyrase-DNA cleavage complexes, subsequently triggering the SOS response, as has been documented in other species like *Mtb*^35^. *Cglu* possesses a functional RecA-LexA regulatory network that is strongly upregulated by mytomycin C, a agent known to cause DNA damage comparable damage to that of FQs^36^. The activation of this network results in a transient block of cell division which significantly amplifies the filamentation of the cells^36^.

In contrast, novobiocin and NBTIs produced milder morphotypes, characterized by moderate elongation and irregular nucleoid structure but without genotoxic stress. While novobiocin primarily inhibits the ATPase activity of GyrB, NBTIs are thought to stabilize a transient gyrase-DNA cleavage complex that blocks the enzyme’s catalytic cycle without generating DSBs. The similarity between these morphotypes, and their close match to those of gyrase-depleted cells, suggests that novobiocin and NBTIs mainly impair DNA relaxation and segregation rather than causing direct DNA damage. Taken together, these findings delineate a spectrum of gyrase associated morphologies in *Cglu*: from moderate topological imbalance (novobiocin), through catalytic inhibition or cleavage-complex stabilization without DSBs (NBTIs and genetic depletion), to combined topological and genotoxic stress (FQs). The close correspondence between chemical and genetic perturbations reinforces the idea that *Cglu* not only serves as a safe and genetically tractable model to study gyrase function *in vivo*, but also provides a powerful cellular readout for discriminating between mechanistic classes of gyrase inhibitors.

Consistent with the above observations, a recent *Msm* study using a different approach, CRISPRi-mediated repression of *gyrA* and *gyrB* combined with quantitative single-cell imaging^37^, reported the same type of morphological changes that we observed under chemical inhibition in *Cglu.* Gyrase depletion in *Msm* caused marked cell elongation and nucleoid disorganization, closely matching the effects of pharmacological gyrase inhibition. Importantly, this study, like ours, distinguished between the outcomes of different inhbitor classes: FQs (e.g. moxifloxacin) induced extreme filamentation and severe nucleoid condensation due to the formation of DSBs and activation of the SOS response, while novobiocin produced milder elongation and polar bulging without pronounced nucleoid loss, consistent with partial inhibition of DNA supercoiling. This convergence, despite distinct methodologies, confirms that disruption of DNA gyrase universally triggers a characteristic morphological signature whose severity reflects the specific mode of enzyme inhibition.

Finally, although the GyrA*_Mtb_* subunit fully restores gyrase activity in the *Cglu gyrA*-depleted strain, the complemented strain displays minor morphological differences. This indicates that, while the enzyme’s core catalytic mechanism is conserved, optimal gyrase function depends on species-specific subunit interplay, as also noted in the cross-species gyrase studies of *Weidlich et al.*^38^. The observed reduction in *in vitro* supercoiling activity might stem from structural mismatch between the GyrA*_Mtb_* DEEE-loop and the significantly longer C-loop of GyrB*_Cglu_* (45 versus 32 residues) (Supplementary Figure 4) ^17,38^. In addition, the C-tail of GyrA*_Mtb_* is markedly shorter than that of GyrA*_Cglu_*, a feature that may further weaken the communication interface with GyrB*_Cglu_* (Supplementary Figure 4)^18^. This disparity could probably disrupt the precise interdomain coupling required to efficiently coordinate ATP hydrolysis with DNA strand passage, resulting in a “functional but less efficient” heterologous enzyme. Nevertheless, the heterologous enzyme is sufficient to support viability despite its reduced efficiency, the shift in phenotype likely reflects a modification of the DNA supercoiling “setpoint”^39^. Because DNA topology directly affects gene expression, any deviation from the optimal superhelical density caused by the heterologous enzyme would alter the bacterial transcriptional profile, resulting in the observed phenotypic differences^39^.

In summary, our findings validate *Cglu* as a reliable, non-pathogenic model for studying DNA gyrase function and inhibition in Corynebacteriales. The strong structural and functional conservation between *Cglu* and *Mtb* gyrases, together with the clear phenotypic readouts described here, underscore the potential of *Cglu* as a surrogate platform for drug discovery, providing a practical and biosafe route to accelerate the identification and characterization not only of new gyrase-targeting compounds but also of new compounds directed against other antibacterial targets.

### Methods

#### Bacterial strains, plasmids and growth conditions

All strains and plasmids used are listed in Supplementary Table 2. *E. coli* DH5α (NEB) or CopyCutter EPI400 (Lucigen) were used for cloning and grown in Luria-Bertani (LB) broth or agar plates at 37 °C, supplemented with 50 μg.mL^-1^ kanamycin when required. For protein production, *E. coli* BL21 (DE3) was grown in 2YT broth supplemented with auto-induction medium (0.5% glycerol, 0.05% glucose, 0.2% lactose) and 50 μg.mL^-1^ kanamycin^40^. *Cglu* ATCC13032 was used as the wild-type strain. *Cglu* strains were grown in controlled CGXII medium at 30 °C and 120 r.p.m., supplemented with 50 μg.mL^-1^ kanamycin when required. For overexpression, CGXII containing 4% sucrose was supplemented with 1% gluconate. For inhibition of the expression CGXII containing 4% sucrose was supplemented with 1% *myo*-inositol.

#### Cloning for recombinant protein production in *E. coli*

The genes encoding for *Cglu* DNA gyrase (GyrA and GyrB) subunits were amplified by PCR from the chromosomal DNA of *Cglu* ATCC 13032 and cloned by Gibson assembly (NEB) into a pT7 vector containing an N-terminal 6-His-SUMO tag upstream of a SUMO cleavage site. To simplify structural studies of the heterotetrameric GyrB_2_GyrA_2_ complex, *Cglu* GyrB and GyrA were fused with a three-amino acid G-D-L linker, following a strategy previously used for *Thermus thermophilus* and *Mycobacterium tuberculosis*^29,17^. DNA gyrase and cloned into a pT7 vector producing a GyrB-GyrA fusion protein (GyrBA) with an N-terminal 6-His-SUMO tag upstream of a SUMO cleavage site. All the oligonucleotides used for PCR amplification of gyrase genes are listed in Supplementary Table 2. All the technical data concerning the *Mtb* DNA gyrase are described in our previous work^19^.

#### *gyrA* and *gyrB* conditional mutants

To obtain the conditional depletion of the gyrase, the endogenous *gyrA* and *gyrB* were individually uncoupled to their native promoter by a terminator and placed under the control of a repressible promoter. Using the two-step recombination strategy with the pk19mobsacB plasmid, we inserted a terminator followed by the native promoter of the inositol phosphate synthase gene (*ino1*), which can be repressed in the presence of *myo*-inositol. The terminator and the *ino1* promoter were amplified by PCR from the pK19-P3323-lcpA plasmid. The 500 bp up-stream region and down-stream region of *gyrA* and *gyrB* were amplified from the chromosomal DNA of *Cglu* ATCC13032. The different fragments were assembled in a pk19mobsacB backbone by Gibson assembly (NEB). The plasmid pK19mobsacB-*P_ino_*-GyrA and pK19mobsacB-*P_ino_*-GyrB were sequenced and electroporated into WT *Cglu* ATCC13032. Genomic integration of pK19mobsacB plasmids was selected on kanamycin and positive colonies were sucrose sensitive. The second round of recombination was confirmed by growth on 10% (w/v) sucrose.

The insertion of the terminator and *ino1* promoter was confirmed by colony PCR and Sanger sequencing (Eurofins). All the oligonucleotides used to obtain and check both *gyrA* and *gyrB* conditional mutants are listed in Supplementary Table 2.

The complementation of the *P_ino_*-*gyrA Cglu* strain was performed using an expression system based on the pTGR shuttle vector, which is compatible with *E. coli* and *Cglu*^25^. In this study, we used a modified version of this vector derived from pTGR5 containing a transcriptional cassette controlled by the inducible *P_gntK_* promoter. This promoter, derived from the *gntK* gene of *Cglu* involved in gluconate metabolism, enables tight and tunable regulation of gene expression, with basal activity in sucrose and strong induction in gluconate. For this purpose, the *gyrA* gene of *Cglu* fused at the C-terminus to mNeonGreen (*gyrA-mNeonGreen*), as well as the *gyrA* gene of *Mtb* (*gyrA_Mtb_*), were cloned under the control of the *P_gntK_* promoter in the pTGR5-derived expression system. The resulting plasmids, named pUMS420 (*P_gntK_*-*gyrA-mNeonGreen*) and pUMS-383 (*P_gntK_*-*gyrA_Mtb_*), were verified by Sanger sequencing (Eurofins) and subsequently electroporated into the *P_ino_*-*gyrA Cglu* strain for complementation experiments. Transformants were selected on controlled medium CGXII supplemented with 4% sucrose agar containing 50 μg.mL^-1^ kanamycin. All the oligonucleotides used to obtain and check this strain are listed in Supplementary Table 2.

#### Protein production and purification

*Cglu* GyrA and GyrB DNA gyrase subunits were produced and purified separately in a three-step process with two affinity chromatography (IMAC and Heparin) followed by a size-exclusion chromatography (SEC) as we previously reported for *Mtb* gyrase subunits^19^.

#### Antibody production and purification

Anti-GyrA*_Cglu_*, anti-GyrA*_Mtb_* and anti-GyrB*_Cglu_* antibodies were raised in rabbits (Covalab) against purified GyrA*_Cg_*, GyrA*_Mtb_* or GyrB*_Cglu_* antigens. For antibody purification, both sera from day 67 post inoculation were purified using a 1 mL HiTrap NHS-activated HP column (Cytiva) loaded with the corresponding antigen according to manufacturer instructions. Sera were diluted in binding buffer (20 mM sodium phosphate (pH 7.4), 500 mM NaCl), loaded onto the column and washed with 7 mL of binding buffer. Antibodies were eluted with 10 mL of elution buffer (100 mM glycine (pH 3), 500 mM NaCl) and neutralized with 1 M Tris (pH 9). Purified antibodies were concentrated to 8 mg.ml^-1^ and mixed 1:1 with 100% glycerol, aliquoted and stored at -20 °C.

#### Western blots

For cell extracts preparation, pellets were resuspended in lysis buffer (50 mM Bis-Tris (pH 7.4), 75 mM 6-aminocaproic acid, 1 mM MgSO4, benzonase and protease inhibitor) and disrupted at 4 °C with 0.1 mm glass beads and using a PRECELLYS 24 homogenizer. Total extracts (40 μg) were run on an SDS–PAGE gel, transferred onto a 0.2 μm nitrocellulose membrane and incubated overnight with a blocking buffer (5% skimmed milk, 1X TBS-Tween buffer) at room temperature. Blocked membranes were incubated for 1h at room temperature, with the corresponding primary antibody diluted to the appropriate concentration in blocking buffer. After washing in TBS-Tween buffer, membranes were probed with an anti-rabbit horseradish peroxidase-linked secondary antibody (Cytiva) for 45 min. For chemiluminescence detection, membranes were washed with 1X TBS-T and revealed with HRP substrate (Immobilon Forte, Millipore). Images were acquired using the ChemiDoc MP imaging system (BioRad). All uncropped blots are shown in Extended data Figure 1. Dilutions use anti-GA-*Cglu* (1:2000), anti-GB-*Cglu* (1:2000), anti-GA-*Mtb* (1:1000), anti-mNeonGreen (1:1000), and anti-rabbit secondary Abs (1:10000).

Western blot data were normalized using the *Total Protein Normalization* approach (Bio-Rad) based on Stain-Free technology. Following gel activation and protein transfer, total lane protein was visualized and quantified with the Image Lab software (Bio-Rad). The total protein signal of each lane was used as a loading control, and the intensity of target bands was normalized to the corresponding total protein signal, thereby minimizing variability due to unequal loading or transfer efficiency.

#### Phase contrast, fluorescence microscopy and image analysis

For antibiotic treatment imaging, a culture of *Cglu* ATCC13032 was grown overnight at 30°C and 120r.p.m in controlled medium CGXII supplemented with 4% sucrose and diluted to OD_600_=1 the following day and grown at 30°C and 120r.p.m until the early exponential phase (5h) before adding the antibiotic to the cultures at a concentration equivalent to 10x MIC value for each antibiotic^28^. After 24h of growth, the cultures were collected for imaging. Cultures (100 μl) were pelleted at 5,200×g washed with fresh medium and diluted to an OD_600_ of 3 for imaging.

For imaging the *P_ino_* strains, cultures were grown in controlled medium CGXII supplemented with 4% sucrose and kanamycin for the complemented strain (50 μg.ml^-1^) at 30 °C with shaking at 120 r.p.m for overnight growth. The following day, cultures were diluted to an OD_600_=1 in CGXII and 4% sucrose (± 1% *myo*-inositol & 1% gluconate), grown at 30°C and 120 r.p.m shaking until the exponential phase (6h), diluted at OD_600_=1 (± 1% *myo*-inositol & 1% gluconate) and grown at 30°C and 120 r.p.m until the following day. Cultures (100 μl) were pelleted at 5,200 × g washed with fresh medium and diluted to an OD_600_ of 3 for imaging.

For membrane and DNA staining, Nile Red (Enzo Life Sciences) and Hoechst 33342 (Thermofisher Scientific) were added to the culture (1.6 μg. ml^−1^ and 2 μg. ml^−1^ final concentration respectively) prior to placing them on 2% agarose pads prepared with controlled medium CGXII. Cells were visualized using a Zeiss Axio Observer Z1 microscope fitted with an Orca Flash 4 V2 sCMOS camera (Hamamatsu) and a Pln-Apo 63X/1.4 oil Ph3 objective. Images were collected with Zen Blue 2.6 (Zeiss). They were segmented using a custom trained version of Omnipose, the software Fiji 70^41^ and the plugin MicrobeJ^42^ version 5.13p to generate violin plots. The experiments were performed as biological triplicates. For statistical analysis, due to the important number of cells analyzed in each sample, Cohen’s *d* value was used to describe effect sizes between different strains independently of sample size:

For antibiotics treatment, values were interpreted according to the intervals of reference suggested by Cohen and expanded by Sawilowsky, as follows: small (n.s.), *d* < 0.50; medium (*), 0.50 < *d* < 0.80; large (**), 0.80 < *d* <1.20; very large (***), 1.20 < *d* < 2.0; huge (****), *d* > 2.0.

For the *P_ino_* strains, values were interpreted according to the intervals of reference suggested by Cohen and expanded by Sawilowsky, as follows: small (n.s.), *d* < 0.20; medium (*), 0.20 < *d* < 0.50; large (**), 0.50 < *d* <0.80; very large (***), 0.80 < *d* < 1.20; huge (****), *d* > 1.20.

#### Spot assay

Overnight cultures of *Cglu* were grown in CGXII controlled medium supplemented with 4% sucrose and, for the complemented strain, kanamycin (50 µg. ml^−1^) at 30°C with 120 r.p.m shaking. The next day, cultures were diluted to an OD_600_ of 1 in CGXII containing 4% sucrose (with or without 1% *myo*-inositol and 1% gluconate) and grown at 30°C with 120 r.p.m shaking to exponential phase (6 h). Cultures were then diluted again to an OD_600_ of 1 and incubated at 30°C until the following day. Derivative strains were normalized to an optical density OD_600_ of 0.5, serially diluted and spotted (10 μl) into medium CGXII supplemented with 4% sucrose with and without 1% of *myo*-inositol, and 1% of gluconate. Plates were incubated for 24h at 30 °C and imaged using a ChemiDoc imaging system (BioRad).

#### DNA gyrase activity assays

Relaxed pBR322 DNA and kinetoplast catenated DNA (kDNA) were purchased from Inspiralis. Supercoiled pB322 DNA was purchased from New England Biolabs. GelRed Nucleic Acid Stain was purchased from INTERCHIM. All other chemicals were purchased for Sigma-Aldrich.

To evaluate the supercoiling activity, reactions were carried out in the presence of 300 ng relaxed pBR322 plasmid DNA and 1 mM ATP in kinetic buffer K1 composed of 40 mM Tris-HCl pH 7.5, 25 mM KCl, 2 mM spermidine, 4 mM DTT, 6 mM magnesium acetate and 100 mM potassium glutamate. Reactions were started by addition of equimolar concentrations (0 to 250 nM) of GyrA/GyrB for WT forms. Heterologous gyrase complexes were assembled by incubating GyrB with a large amount of the GyrA subunit from the other species (220 nM) for 3 minutes at 37°C prior to the reaction. After one hour at 37°C, samples were then supplemented with loading dye (50% glycerol and 0.025% bromophenol blue) and were analyzed by migration on 1% agarose gel in 1X TAE Buffer. Bands were visualized by UV illumination after poststaining with GelRed Nucleic Acid Stain.

To evaluate decatenation activity, reactions were similarly conducted in the presence of 250 ng kDNA and 1 mM ATP in kinetic buffer K2 composed of 40 mM Tris-HCl pH7.5, 10 mM NaCl, 10 mM DTT, 6 mM magnesium acetate and 250 mM potassium glutamate. Reactions were started by addition of 300 nM of each subunit and incubated for one hour at 37°C. Then, reaction mixtures were treated with 0.1 mg.mL^-1^ of proteinase K and 0.2% SDS and incubated for one hour at 37°C. After addition of loading dye, samples were then analysed as for supercoiling assays. All data was replicated three independent times.

#### Cryo-electron microscopy studies

##### Nucleic acid preparation

A 150 bp DNA duplex was reconstituted using two phosphorylated asymmetric synthetic oligonucleotides obtained from Eurogentec. Oligonucleotides ME-73b, ME-77b and their corresponding complementary strands (Supplementary Table 2) were dissolved in DNase-free water at 1 mM concentration. The 150bp double stranded DNA was assembled by mixing at 1:1 molar ratio for each oligonucleotide, annealed by incubating at 95°C for 2 min and then decreasing the temperature by 1°C every 1 min until reaching 20°C.

##### Nucleoprotein complex formation for cryo-EM

The purified *Cglu* GyrBA fusion protein was mixed with the 150 bp dsDNA at 1:1 molar ratio with a final concentration of 2 µM. GyrBA, DNA and moxifloxacin mixtures at 40 µM were incubated for 10 min at 37°C. 5’-(β,γ-imido)triphosphate (Sigma) was then added to the ternary complex at 25 µM and further incubated for 30 min at 30°C. The complex was stored at 4°C until sample freezing on cryo-EM grids.

##### Cryo-EM sample preparation

AuF R0.6/1.0 300 mesh grids were glow-discharged using a Solarus II plasmacleaner (Gatan, Inc), for 10 sec at 5 W prior to the application of 4 µL of the complex. The grids were plunge-frozen in liquid ethane using a Vitrobot Mk-IV (Thermo Fisher Scientific) set at 8°C and 100% humidity, blot time 3 s and blot force + 20.

##### Cryo-EM data collection

Cryo-EM imaging was performed on a Titan Krios3 G4 microscope (FEI) (EMBL, Heidelberg) operated at 300 kV equipped with a Falcon4i camera (Gatan) and a GIF Quantum energy filter (Gatan). The images were recorded in EFTEM nanoprobe mode with Serial EM 4.2.0beta in super-resolution counting mode with a pixel size of 0.731 Å per pixel, with a defocus range between -2.0 and -0.7 μm. Two datasets with tilt angle of 0° and 30° were collected with a total dose of 50 e^-^/Å^2^ distributed on 40 frames. Statistics for cryo-EM data collection for the two datasets are listed in Supplementary Table 2.

##### Cryo-EM and image processing

All image processing steps were done using CryoSPARC^43^. Movies motion correction was performed using *Patch Motion Correction*. The contrast transfer function (CTF) parameters were estimated using *Patch CTF estimation*. A first round of particle picking done with the *blob picker* tool was used to generate 2D classes that were subsequently used as templates for a second round of template-based particle picking. After several rounds of 2D classification, the particles from classes displaying high-resolution features were selected and used to generate reference-free 3D *ab initio* models. The particles were further classified in 3D using *Heterogenous Refinement and 3D Classification.* After particles classification, Non-Uniform (NU) refinement was performed, resulting in a final map at ∼3.2 Å resolution (Supplementary Figure 17).

##### Model building and refinement

An initial model was built by fitting an AF3 model of *Cglu* gyrase cleavage core into the cryo-EM map. The double-stranded DNA was traced and moxifloxacin molecules were placed into the corresponding density using using Coot^44^. Subsequently, several cycles of real-space refinement were done using Phenix^45,46^.

## Acknowledgments

We gratefully acknowledge the C2RT core facilities at the Institut Pasteur, P. England, B. Raynal, S. Brûlé (PFBMI), J. Fernandes and A. Salles (UtechS PBI / Imagopole, supported by France BioImaging; ANR-10–INSB–04; Investments for the Future). The Nanoimaging Core at Institut Pasteur is acknowledged for support with image acquisition. The Nanoimagine Core was created with the help of a grant from the french Government’s Investissements d’Avenir program (EQUIPEX CACSICE – Centre d’analyse des systèmes complexes dans les environnements complexes, ANR-11-EQPX-0008). This work was supported by the IdEx Université Paris Cité, ANR-18-IDEX-0001 (S.P.), Fondation pour la Recherche Médicale, FRM, EQU202303016284 (P.M.A.), Institut Pasteur PTR_726_BactImMorph (A.M.W.) and by institutional grants from the Institut Pasteur, the CNRS, and Université Paris Cité. Molecular graphics were done with ChimeraX, developed at UCSF with support from NIH (R01-GM129325) and NIAID. E.Y. and Y.W. acknowledges a PhD fellowship from the Médicament, Toxicologie, Chimie, Imageries (MTCI ED 563), Université Paris Cité. Computational analyses were performed using the SBGrid software environment, which provides curated, version-controlled structural biology applications and reproducible execution environments across platforms^47^. We would like to sincerely thank Meike Baugmart for her valuable advice in the generation of the *P_ino_* strains. We are also grateful to Mariano Martinez and Yves Marie Boudehen for their support with the molecular biology experiments, and to Giacomo Carloni for generating the Clinker diagram. A grateful ackowledgment to Julienne Petit (JP) for her expertise in *Cglu* cellular imaging and analysis, and for her valuable training of Yaëlle Wormser.

## Author contributions

YW, EY, AS, AG, and SP conceptualized and designed experiments; AS conducted genetic experiments: cloning of *gyrA*, *gyrB* and *gyrBA* of *Cglu* for expression in *E. coli,* design of the *P_ino-_gyrA* and *P_ino-_gyrB* protocol, training of EY; EY conducted genetic experiments: *P_ino-_gyrA* and *P_ino-_gyrB Cglu* strains; MBA conducted genetic experiments: cloning of *gyrA-mNeon* for expression in *Cglu;* YW made the WBs for the *P_ino-_gyrB*/*gyrA* strains characterization; YW conducted the Spot tests; EY and EL conducted all the preliminary microbiological tests; YW acquired and analyzed microscopy images after being trained by JP; AG and EC conducted enzymatic activity; EY and AG purified proteins; EY, FG and SP prepared the cryo-EM grids; FG and SP collected the data; AM processed the cryo-EM data; YW, EY and SP analyzed the 3D structures; YW, AG and SP prepared the figures; AG made the AF3 prediction of the heterologous enzymes; SP supervised the studies; AA, PMA, AMW and SP provided scientific and strategic guidance and funding; YW, AG and SP wrote the original draft of the manuscript; All authors revised and edited the paper.

## Supplementary Figures

**Supplementary Figure 1.**
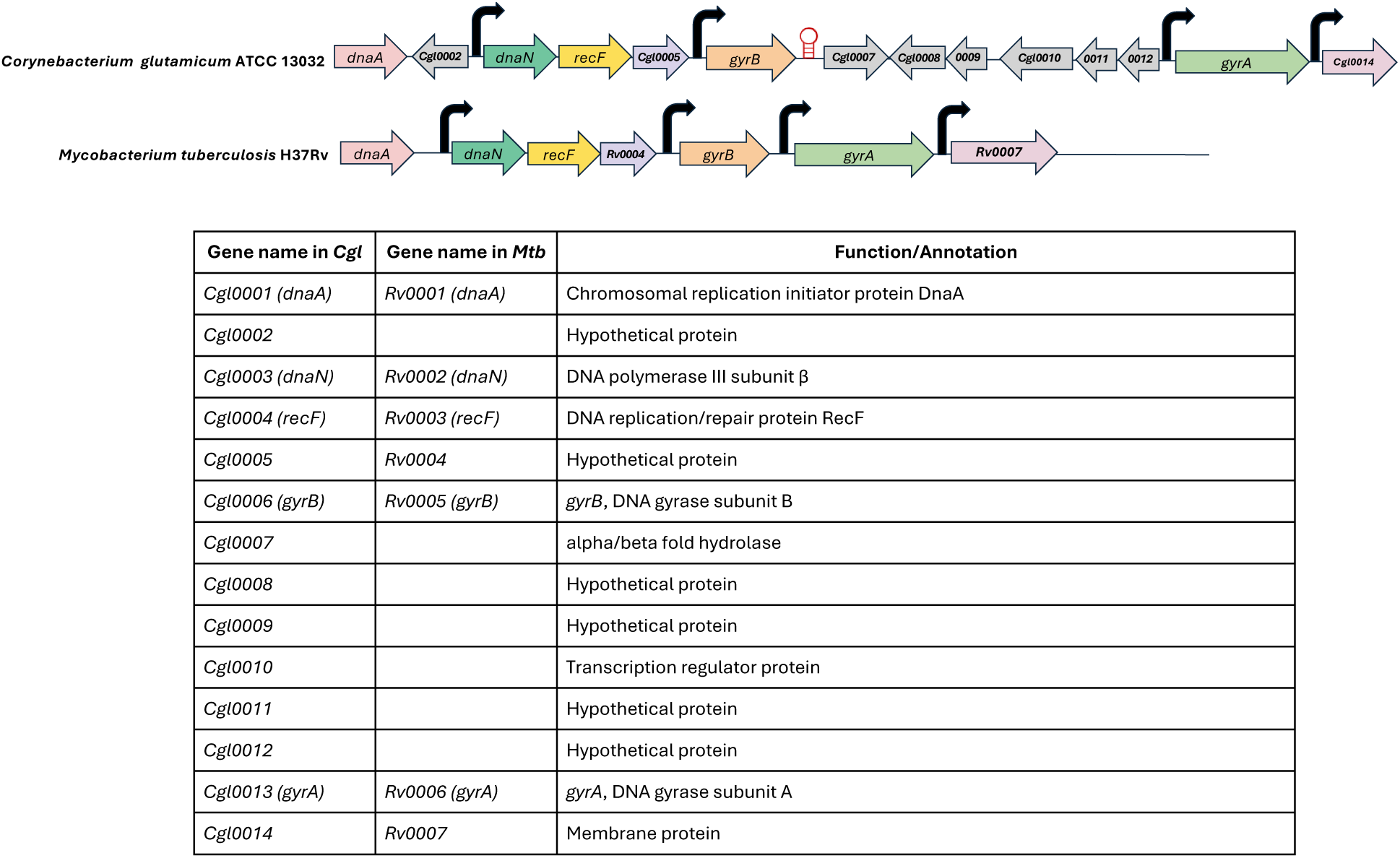
Comparison of the *gyrB*–*gyrA* genomic locus organization in *Cglu* and *Mtb*. Schematic representation of the genomic organization of the *gyrB–gyrA* locus and surrounding genes in *Cglu* ATCC 13032 and *Mtb* H37Rv. Genes belonging to the same operon are indicated with the same color. Functional annotations and gene correspondences (orthologs) between the two species are listed below the maps. Sequence identity percentages between *Cglu* and *Mtb* orthologs were obtained using NCBI BLAST. Genomic organization for *Cglu* and *Mtb* was adapted from *Pfeifer-Sancar et al.*^1^ and *Cole et al.*^2^

**Supplementary Figure 2.**
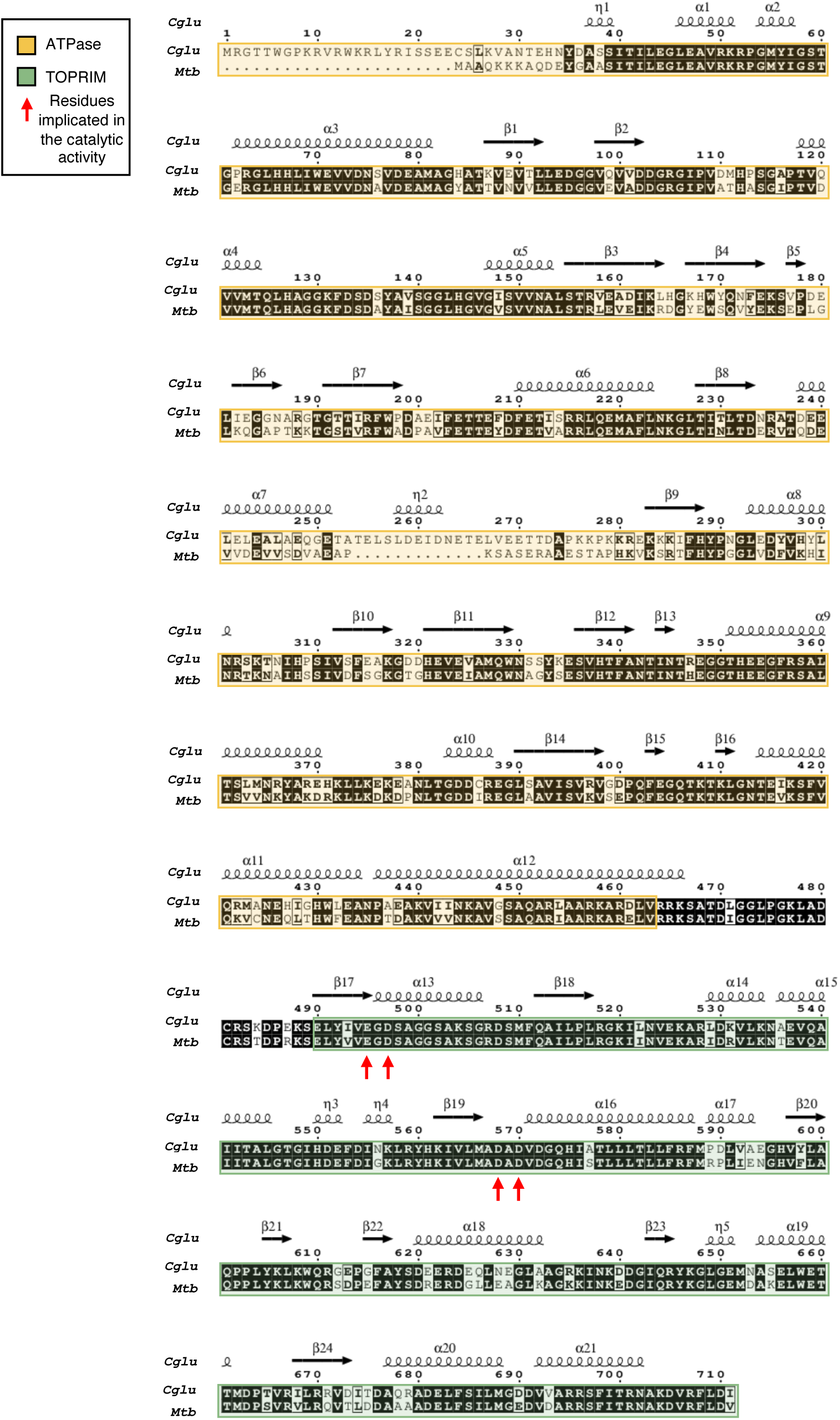
Sequence alignment of GyrB subunits from *Cglu* and *Mtb*. Secondary structure elements predicted for *Cglu* are shown above the alignment. Domains are color-coded as follows: yellow for the ATPase domain, and green for the TOPoisomerase–PRIMase (TOPRIM) domain. Alignment was done with Clustal W 2.0^3^ and ESPript 3.0^4^.

**Supplementary Figure 3.**
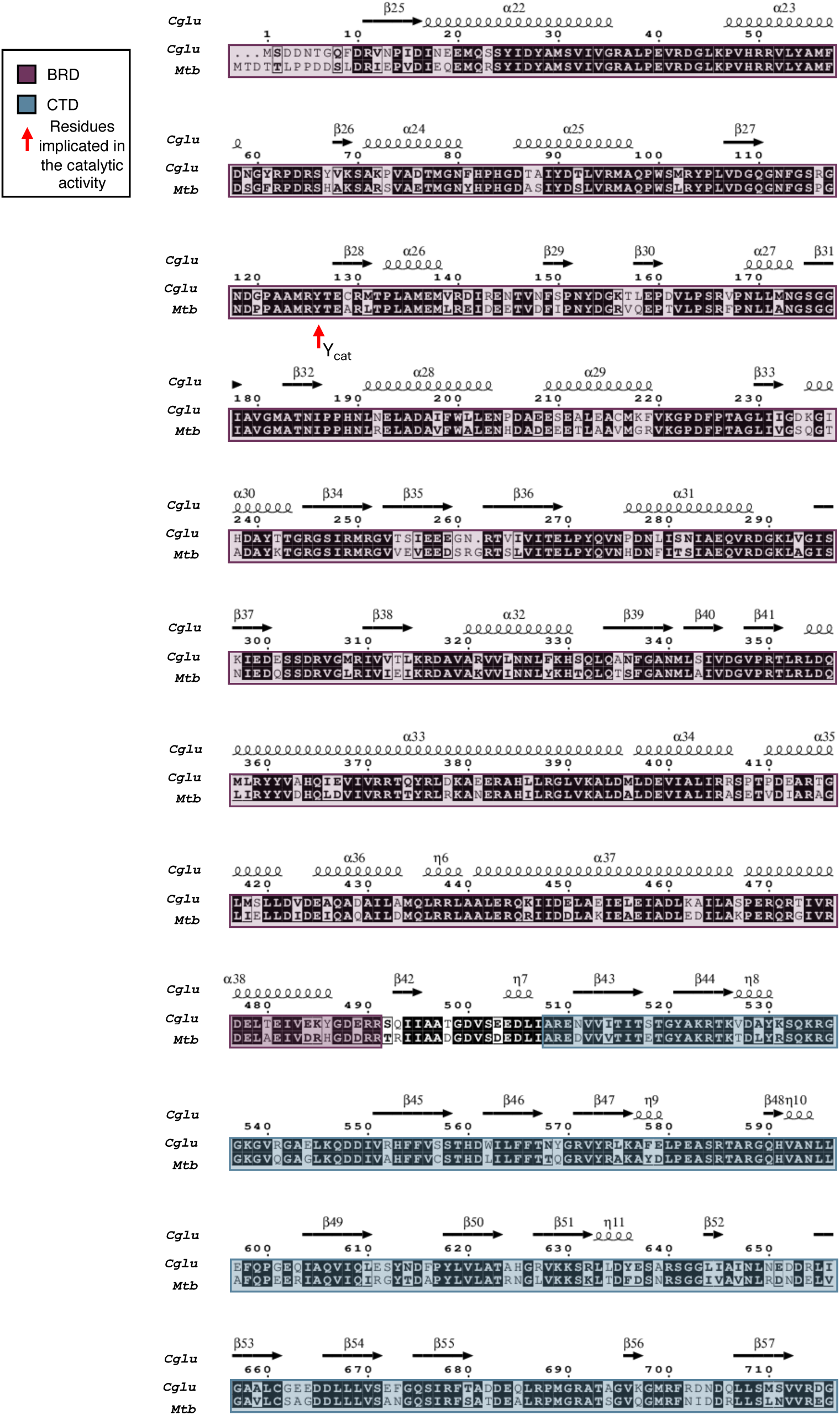
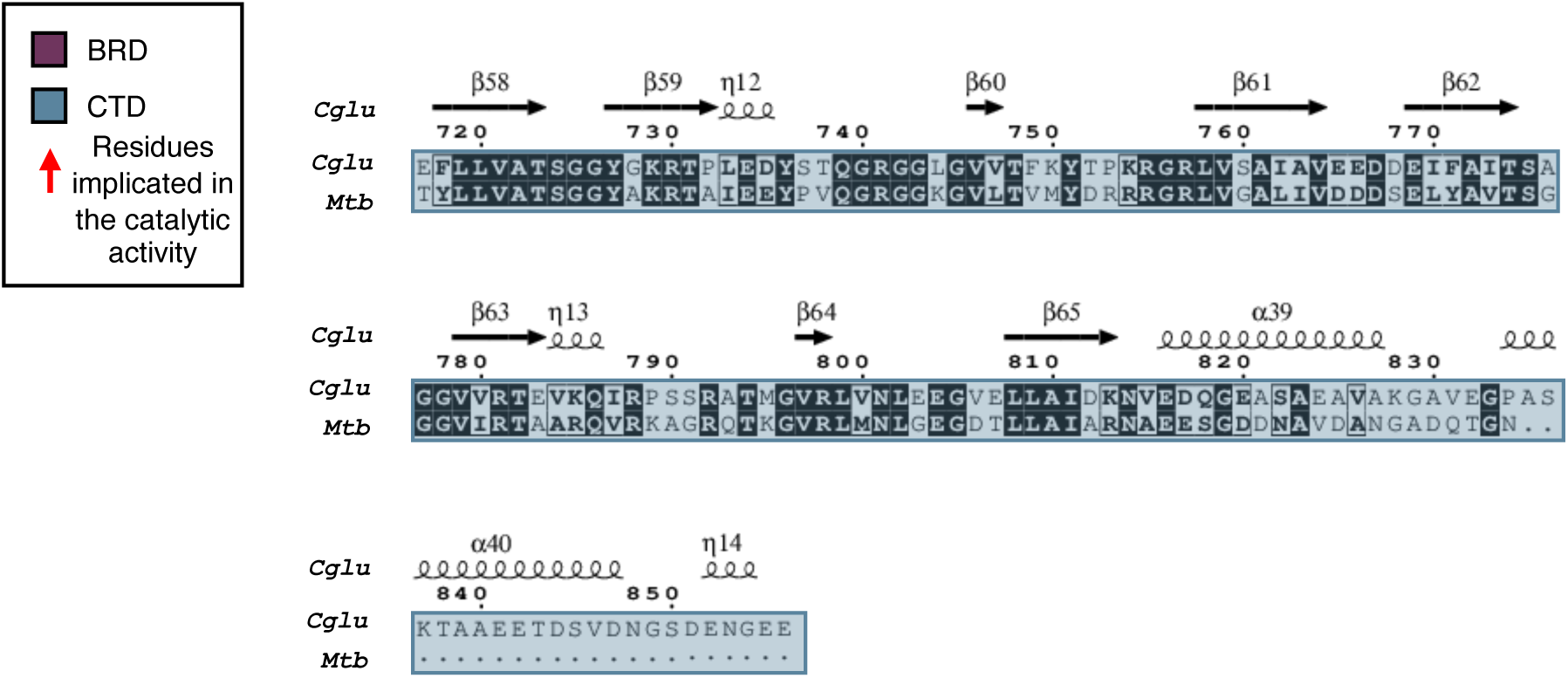
Sequence alignment of GyrA subunits from *Cglu and Mtb*. Secondary structure elements predicted for *Cglu* are shown above the sequence. Domains are color-coded as follows: purple for the the Breakage Reunion Domain (BRD), and blue for the the C-terminal domain (CTD) domain. Alignment was done with Clustal W 2.0^3^ and ESPript 3.0^4^.

**Supplementary Figure 4.**
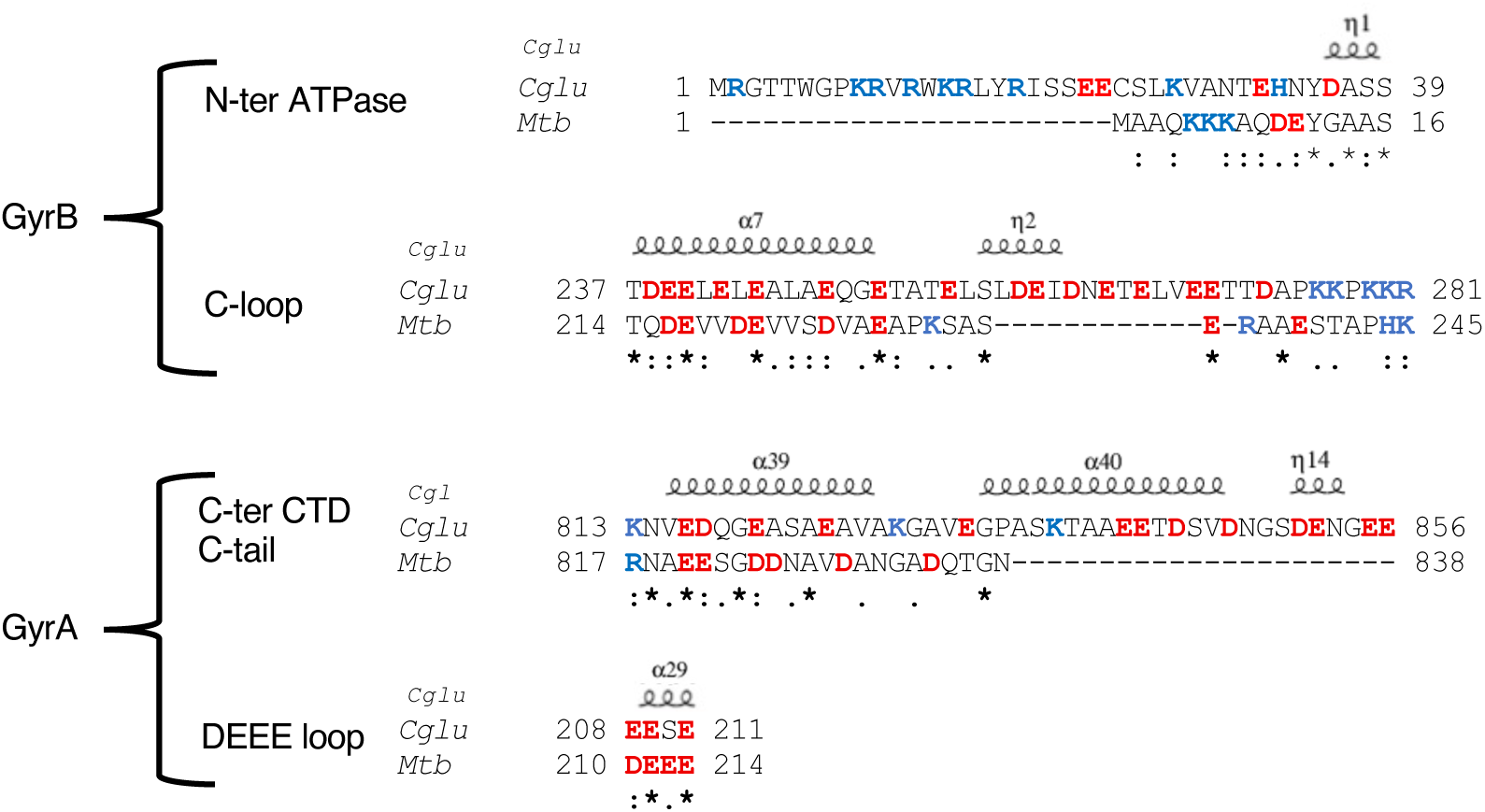
Sequence alignment of the residues from the N-terminal region of GyrB, the C-loop (insertion in the ATPase domain of GyrB) and the C-tail (the extension of the CTD) from *Cglu* and *Mtb* gyrases. Acidic residues are highlighted in red and basic residues in blue. The predicted secondary structures of *Cglu* are indicated above the sequences. In the GyrB subunit, *Cglu* gyrase displays an extended N-terminal region (39 residues in *Cglu* versus 16 in *Mtb*), harboring residues of similar physicochemical properties. The Corynebacteriales*-*specific C-loop (residues 214-245 in *Mtb* / 237-281 in *Cglu*), within the ATPase domain is present in both enzymes but is slightly longer in *Cglu* (45 residues versus 32 in *Mtb*). This motif, located between the last two β-strands of the GHKL domain, exhibits variable amino acid sequences among Corynebacteriales but maintains a conserved acidic composition (about one-third acidic residues in both *Cglu* and *Mtb*). Another notable difference between the two species lies in the extreme C-terminus of the GyrA subunit (the extreme C-terminus of the CTD)^5^. This element referred as the C-tail is highly acidic in both species (the predicted pI is around 4.0; Supplementary Table 1) but is significantly longer in *Cglu* (44 residues for *Cglu* versus 22 for *Mtb* C-tail).

**Supplementary Figure 5.**
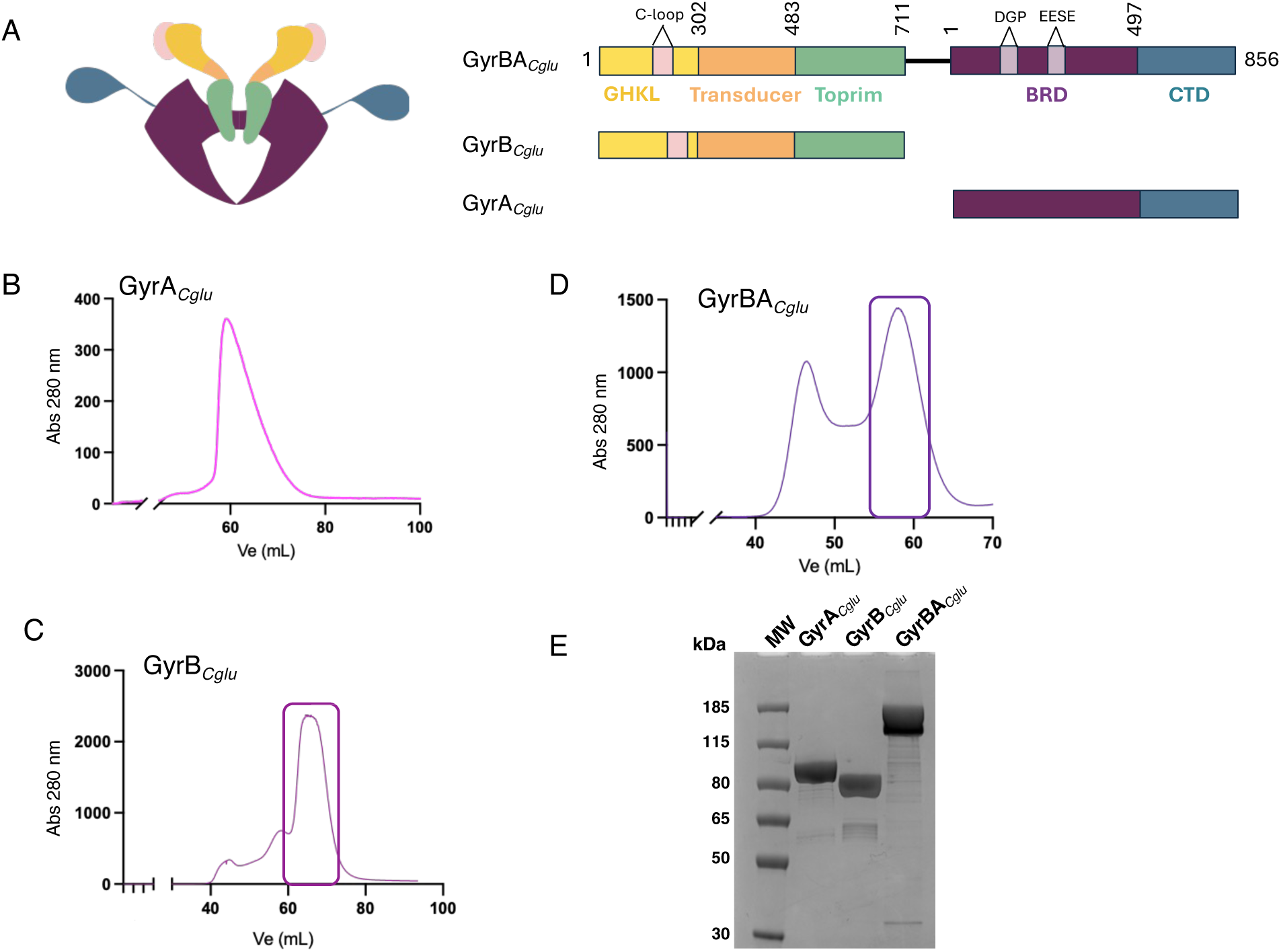
Purification of GyrA, GyrB and GyrBA from *Cglu*. (A) Domain organization of the *Cglu* DNA gyrase constructs used in this study. Functional regions are colored and labeled. The linker connecting the DNA gyrase B and A subunits is represented in a gray dashed line (3-amino-acid linker, G-D-L). The pink insertion in the GHKL domain corresponds to the C-loop (residues 237-281 in GyrB), the light purple insertions in the BRD correspond to the DGP and EESE. These motifs previously described in the BRD domain of *Mtb* gyrase are also present in *Cglu*: the DEEE-loop, exposed on the solvent-accessible surface and extending one of the two α-helices, was shown to contribute to the stabilization of the enzyme in its resting state through interactions with the C-loop^6^. The DPP motif, located in the loop connecting the α3-α4 DNA-binding motif and the catalytic tyrosine residue, participates in DNA binding^7^. Both motifs are conserved in *Cglu*, though with slight variations: the DEEE-loop corresponds to an EESE-loop (residues 211-214 in *Mtb* / 208-211 in *Cglu*), and the DPP motif (residues 122-124 in Mtb / 119-121 in *Cglu*) is replaced by a DGP sequence in *Cglu*. (B-D) Chromatograms from the third purification step of each protein by gel filtration on a Superdex200 (Cytiva Life Sciences) at a flow rate of 1 mL/min. Peaks corresponding to protein of interest are highlighted by a rectangle. (E) SDS-PAGE analysis of purified proteins GyrA, GyrB, and GyrBA of *Cglu*.

**Supplementary Figure 6.**
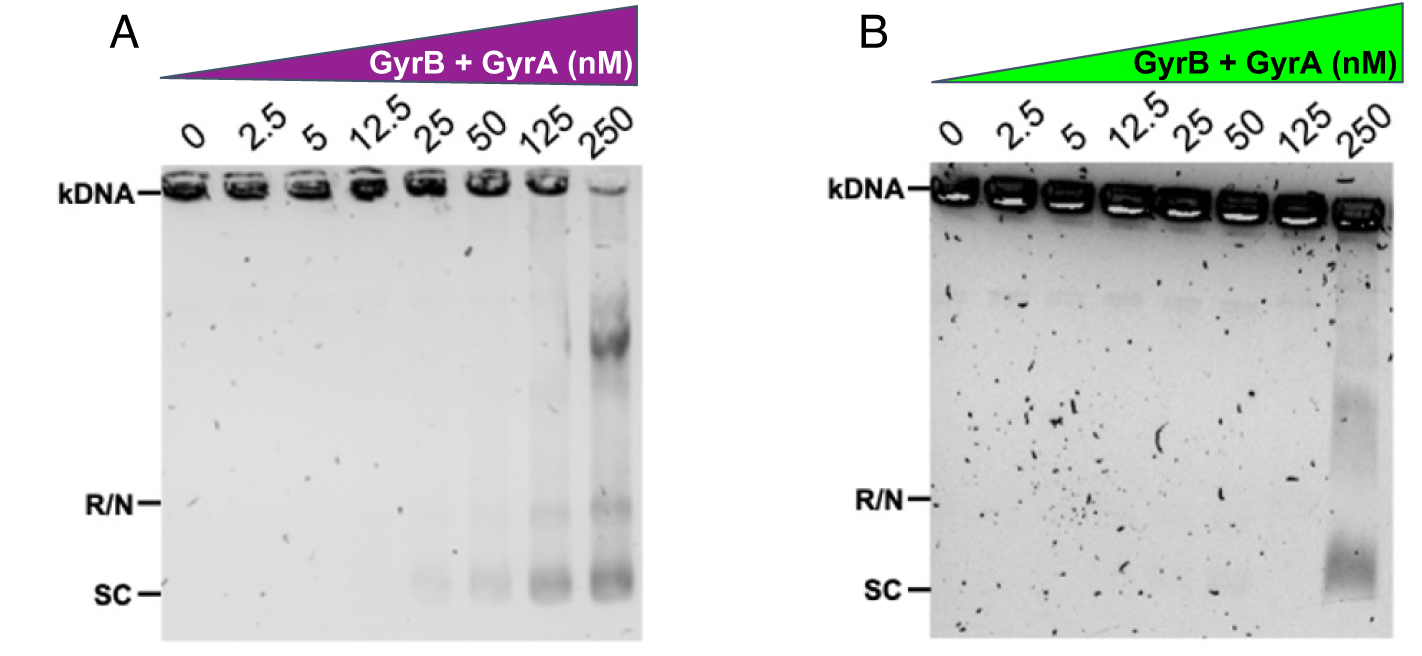
Decatenation activity assay of *Cglu* DNA gyrase. (A) DNA decatenation activity for *Cglu* in the presence of 300 nM of each subunit on variable concentrations of kDNA. (B) DNA decatenation activity for *Mtb* in the presence of 300 nM of each subunit on variable concentrations of kDNA. R, relaxed DNA; N, Nicked DNA; SC, supercoiled DNA; kDNA, kinetoplastic DNA

**Supplementary Figure 7.**
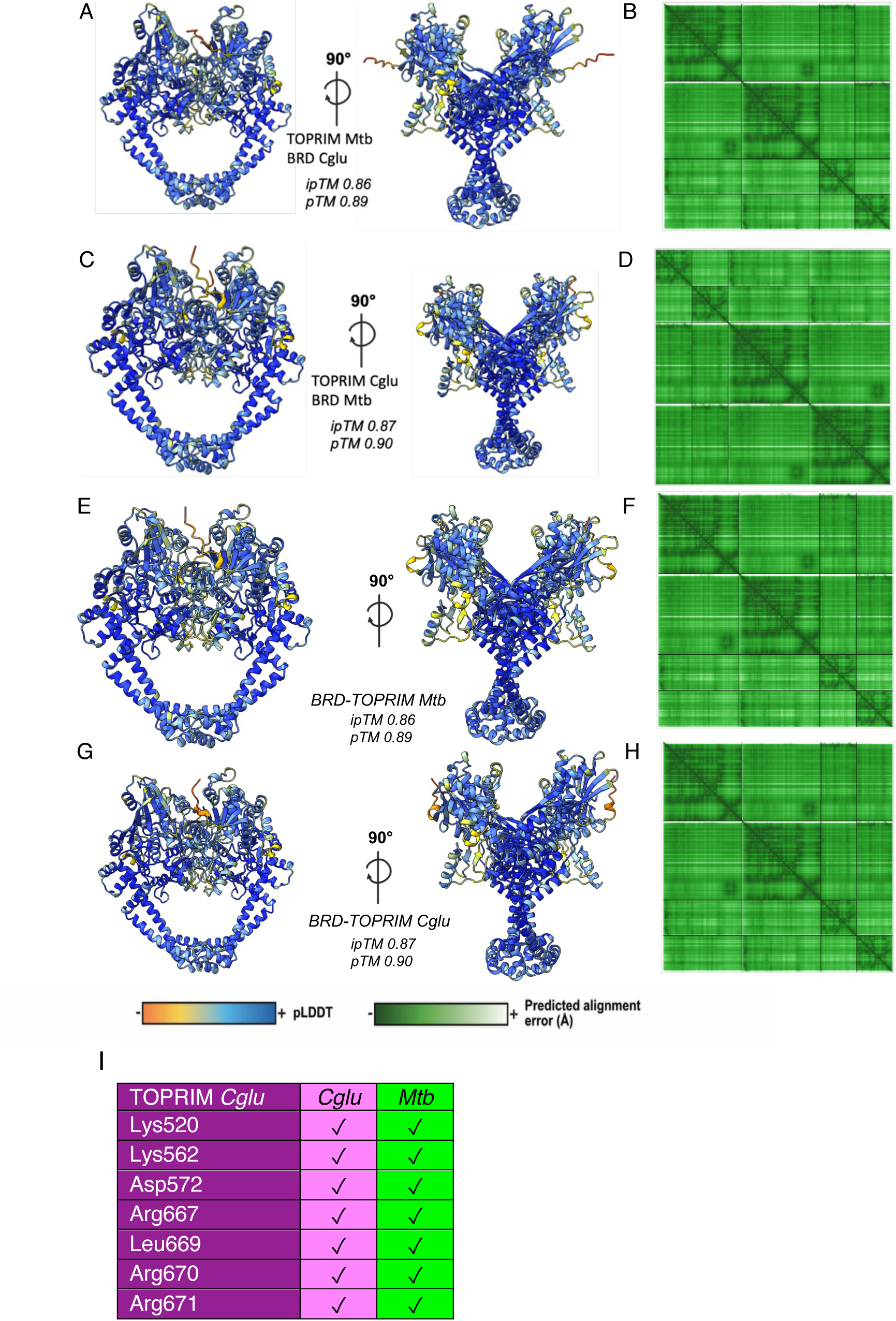
3D structure prediction of the chimeric DNA gyrases. (A, C, E, G) AF prediction for the TOPRIM-*Mtb*/BRD-*Cglu* heterotetramer (ipTM/pTM=0.86/0.89), TOPRIM-*Mtb*/BRD-*Mtb* (ipTM/pTM=0.87/0.90), (ipTM/pTM=0.86/0.89) and (ipTM/pTM=0.87/0.90). The models are color-coded by model confidence. (B, D, F, H) Corresponding residue-residue alignment confidence plots. I, Highly conserved BRD-TOPRIM interaction network in *Cglu* and *Mtb* (See Extended data Tables 1 to 4)

**Supplementary Figure 8.**
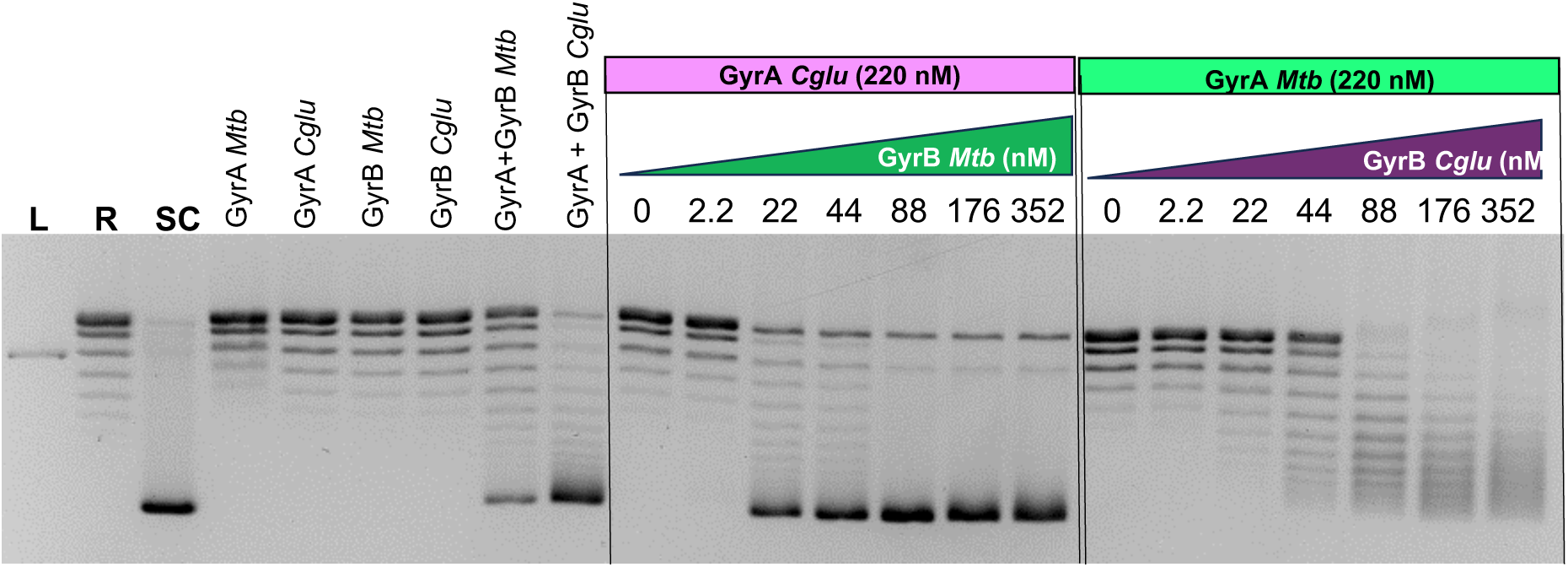
Heterologous DNA gyrases exhibit functional supercoiling activity. Gel of activity assays of negative supercoiled DNA. Supercoiling activity assay on 300 ng as detailed in Methods on relaxed pBR322 plasmid in the presence of 220 nM of GyrA *Cglu* and varying concentrations of GyrB *Mtb* or 220 nM of GyrA *Mtb* in the presence of varying concentrations of GyrB *Cglu* (Shown in Figure 1). Negative controls include the presence of sole proteins in the absence of the other subunit. Positive controls included for *Mtb* gyrase (520 nM GyrA+ 197 nM GyrB) and *Cglu* gyrase (85 nM GyrA + 3 μM GyrB). Mean and standard deviation shown for n = 2.

**Supplementary Figure 9.**
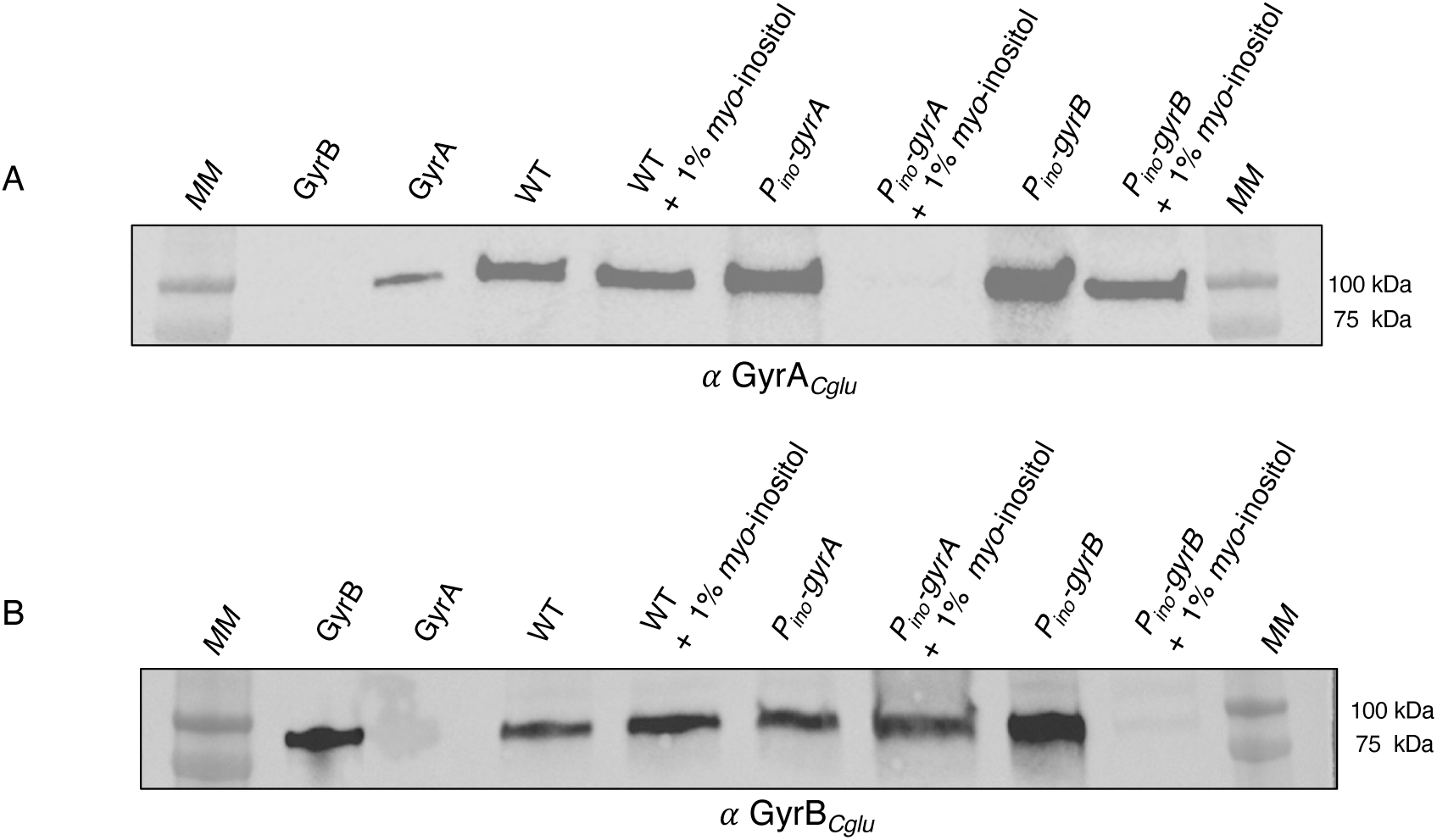
*P_ino_-gyrA* and *P_ino_-gyrB* depletion. Western blots of whole-cell extracts of WT, *P_ino_-gyrA* and *P_ino_-gyrB* strains, grown in either 4% sucrose or 4% sucrose +1% *myo*-inositol (depletion). The blot was revealed with an: (A) anti-GyrA*_Cglu_* antibody. (B) anti-GyrB*_Cglu_* antibody. 8 ng of recombinant GyrA*_Cglu_* and GyrB*_Cglu_* purified protein was deposited on the gel as controls. Molecular weight markers (kDa) are shown on each side of the blot. The blots shown are representative of experiments made independently in triplicate.

**Supplementary Figure 10.**
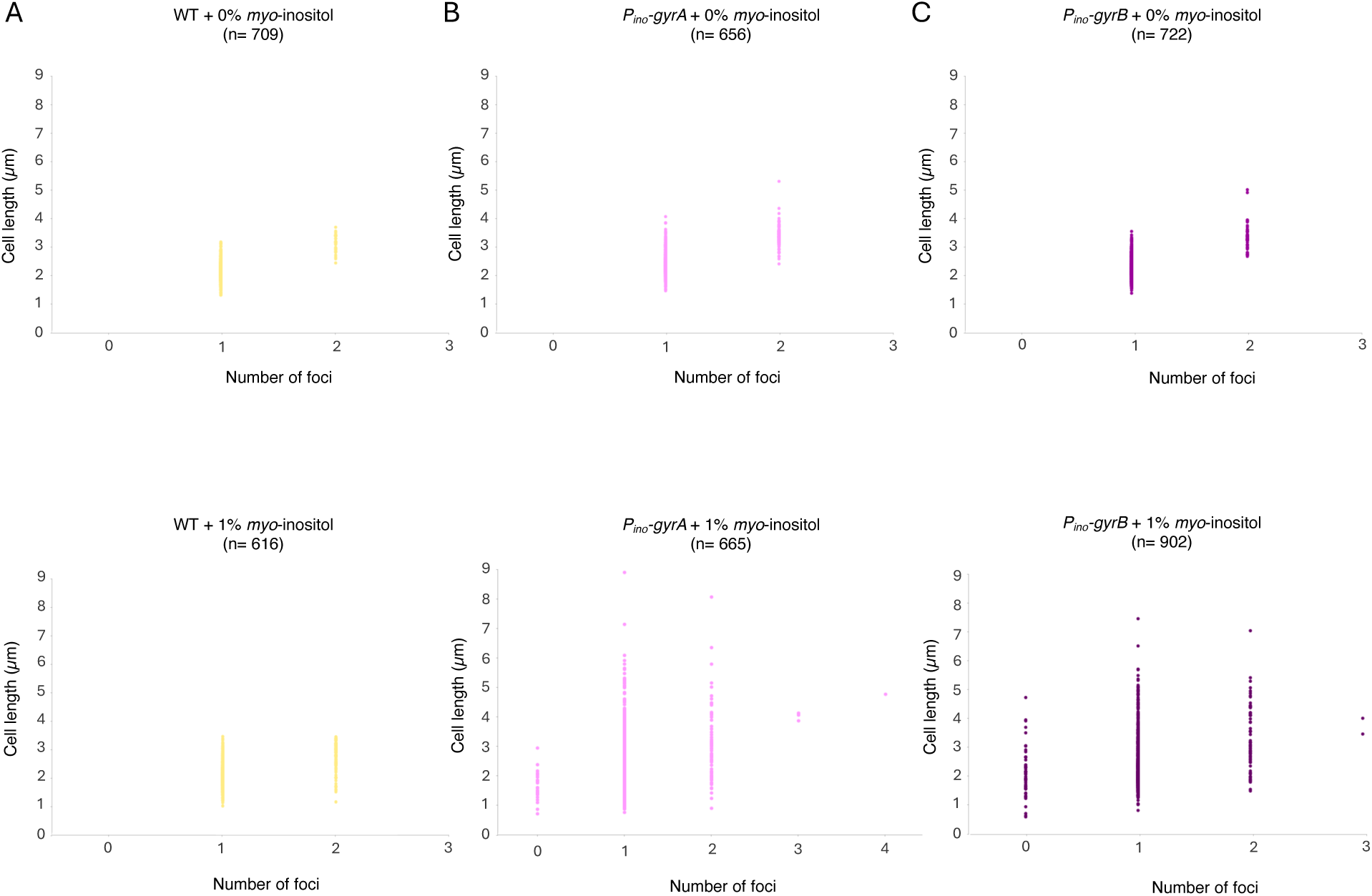
Distribution of cell length according to the number of DNA foci in *Cglu* wild-type and gyrase depletion strains grown with or without *myo*-inositol. Scatter plots show the distribution of cell length as a function of the number of fluorescent foci per cell for the wild-type strain (WT) and the depletion strains *P_ino_-gyrA* and *P_ino_-gyrB* grown at 24 hours with or without *myo*-inositol for cell grown in controlled medium CGXII supplemented with 4% sucrose. Panels A, B, and C correspond to WT, *P_ino_-gyrA*, and *P_ino_-gyrB*, respectively. Each dot represents a single cell, (n) represent the number of cells used in the analyses.

**Supplementary Figure 11.**
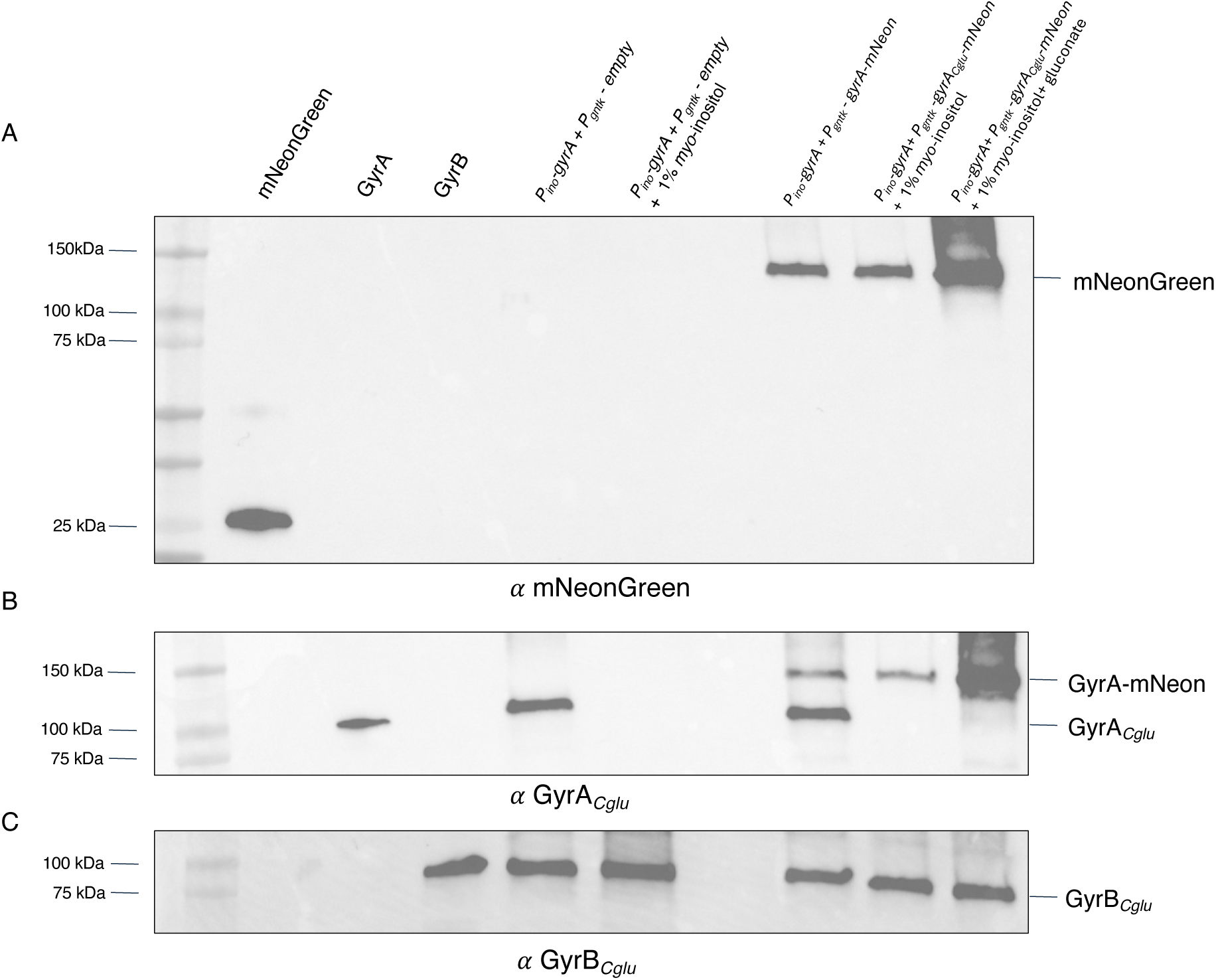
*P_ino_-gyrA* strain complemented with *P_gntk_-gyrA* of *Cglu* fused to mNeonGreen. Western blots of whole-cell extracts of *P_ino_-gyrA* strains complemented with *P_gntK_-empty* (empty plasmid) and *P_gntK_-gyrA_Cglu_-mNeon*, grown in either 4% sucrose, 4% sucrose +1% *myo*-inositol (depletion) or 4% sucrose +1% *myo*-inositol + 1% gluconate (depletion + induction of the expression of the gene controlled by the *P_gntk_* promoteur). The blot was revealed with an: (A) anti-mNeon antibody. (B) anti-GyrA*_Cglu_* antibody. (C) anti-GyrB*_Cglu_* antibody. 8 ng of purified recombinant mNeonGreen, GyrA *Cglu* and GyrB *Cglu* proteins were used as controls for antibody specificity. Molecular weight markers (kDa) are shown on each side of the blot. The blots shown are representative of experiments made independently in triplicate.

**Supplementary Figure 12.**
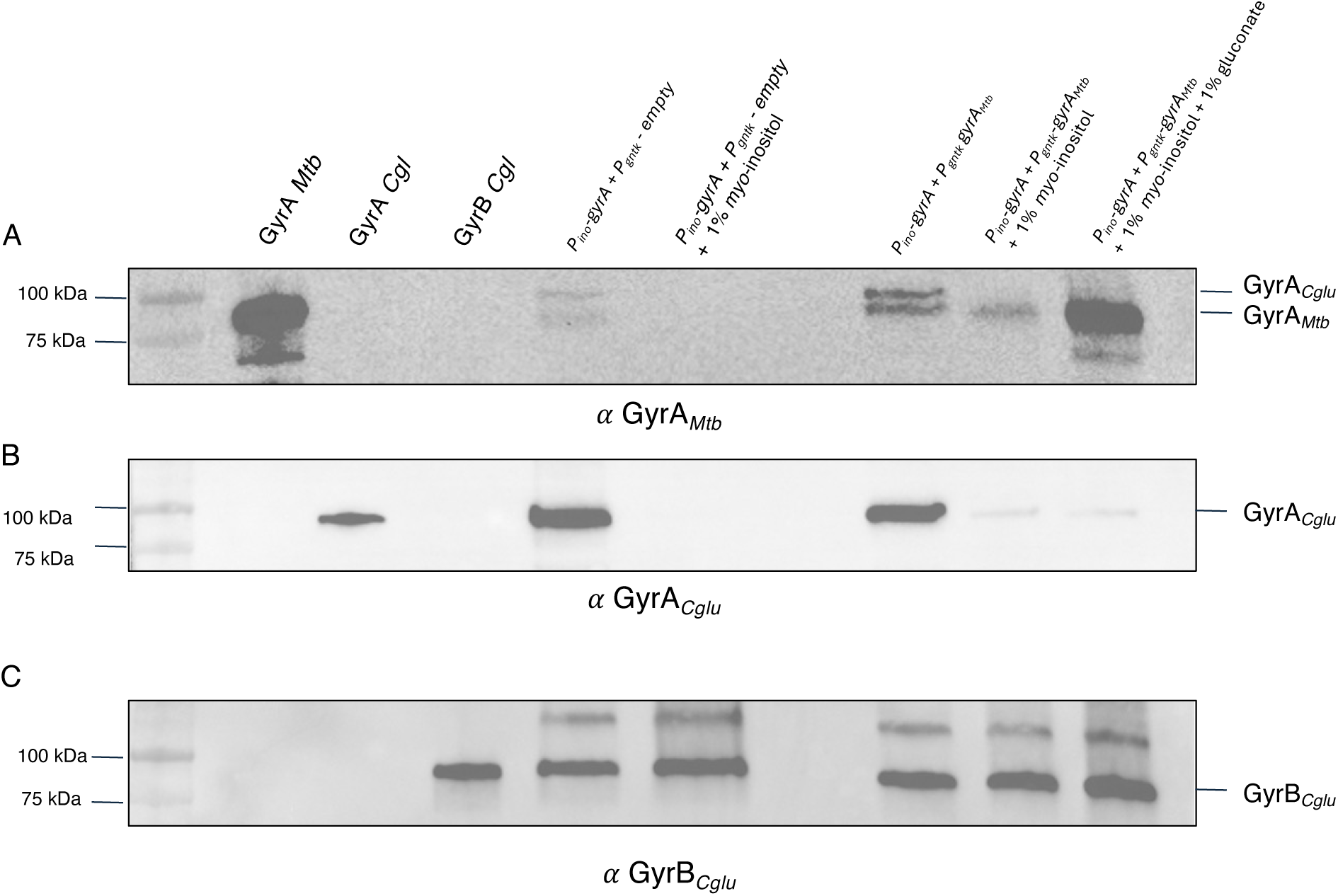
*P_ino_-gyrA* strain complemented with *P_gntk_-gyrA* of *Mtb*. Western blots of whole-cell extracts of *P_ino_-gyrA* strains complemented with *P_gntK_-empty* (empty plasmid) and *P_gntK_-gyrA_Mtb_*, grown in either 4% sucrose, 4% sucrose +1% myo-inositol (depletion) or 4% sucrose +1% myo-inositol + 1% gluconate (depletion + induction of the expression of the gene controlled by the *P_gntk_* promoteur). The blot was revealed with an: (A) anti-GyrA*_Mtb_* antibody. (B) anti-GyrA*_Cglu_* antibody. (C) anti-GyrB*_Cglu_* antibody. 8 ng of purified recombinant GyrA *Mtb,* GyrA *Cglu* and GyrB *Cglu* proteins were used as controls for antibody specificity. Molecular weight markers (kDa) are shown on each side of the blot. The blots shown are representative of experiments made independently in triplicate.

**Supplementary Figure 13.**
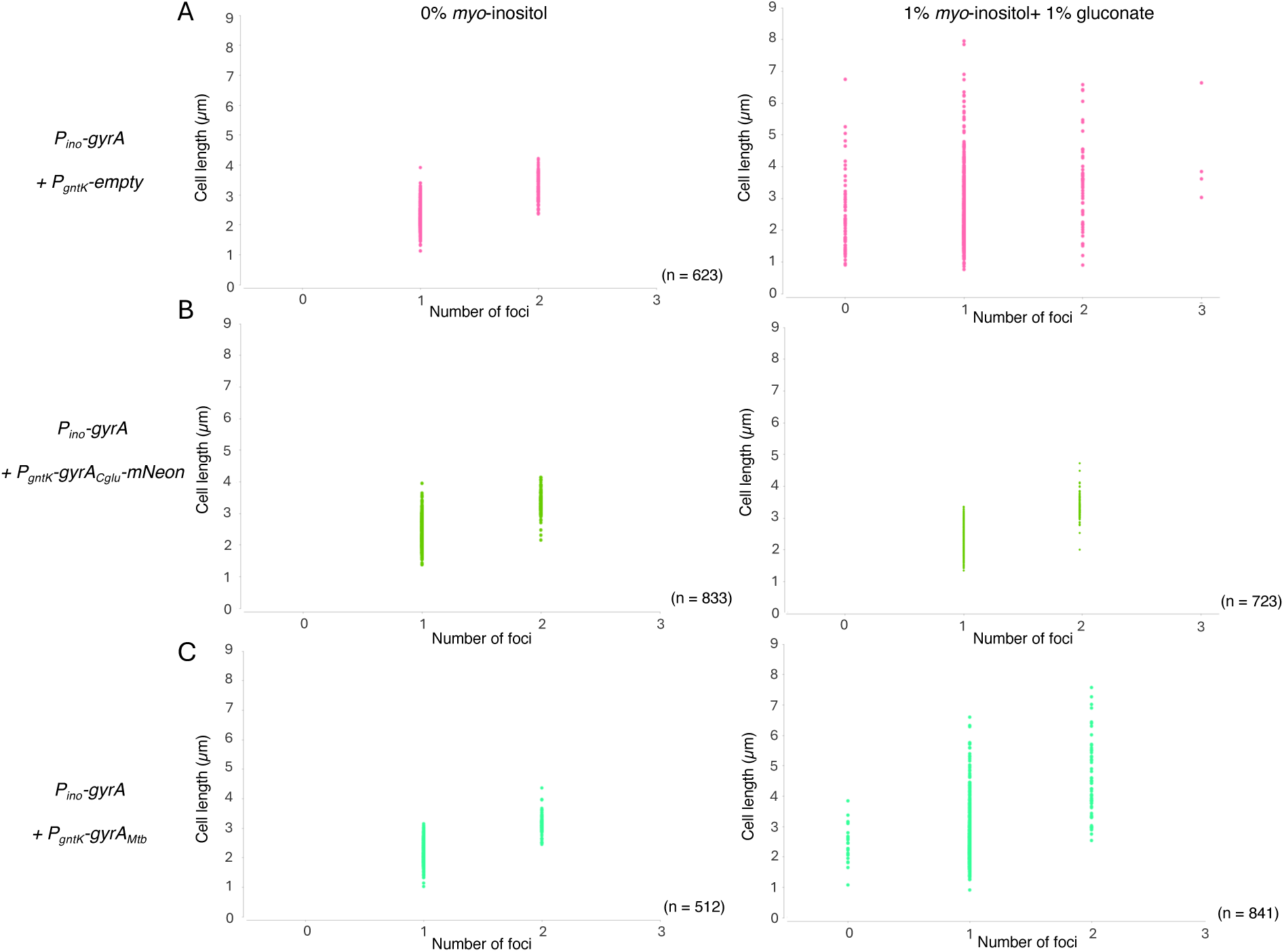
Distribution of cell lengths according to the number of DNA foci in *Cglu* complemented strains. Scatter plots show the distribution of cell lengths relative to the number of fluorescent foci per cell of the depletion strain *P_ino_-gyrA* + *P_gntK_ empty* and the complementation strains *P_ino_-gyrA* + *P_gntK_ gyrA-mNeon* and *P_ino_-gyrA* + *P_gntK_ gyrA_Mtb_* grown at 24 hours with or without *myo*-inositol and gluconate for cell grown in controlled medium CGXII supplemented with 4% sucrose. (A) *P_ino_-gyrA* + *P_gntK-_*empty (negative control), grown at 24 hours with or without *myo*-inositol and gluconate for cell grown in controlled medium CGXII supplemented with 4% sucrose. (B) *P_ino_-gyrA + P_gntK_-gyrA-mNeon*, grown at 24 hours with or without *myo*-inositol and gluconate for cell grown in controlled medium CGXII supplemented with 4% sucrose. (C) *P_ino_-gyrA + P_gntK_-gyrA_Mtb_* grown at 24 hours with or without *myo*-inositol and gluconate for cell grown in controlled medium CGXII supplemented with 4% sucrose. Each dot represents a single cell, (n) represent the number of cells used in the analyses.

**Supplementary Figure 14.**
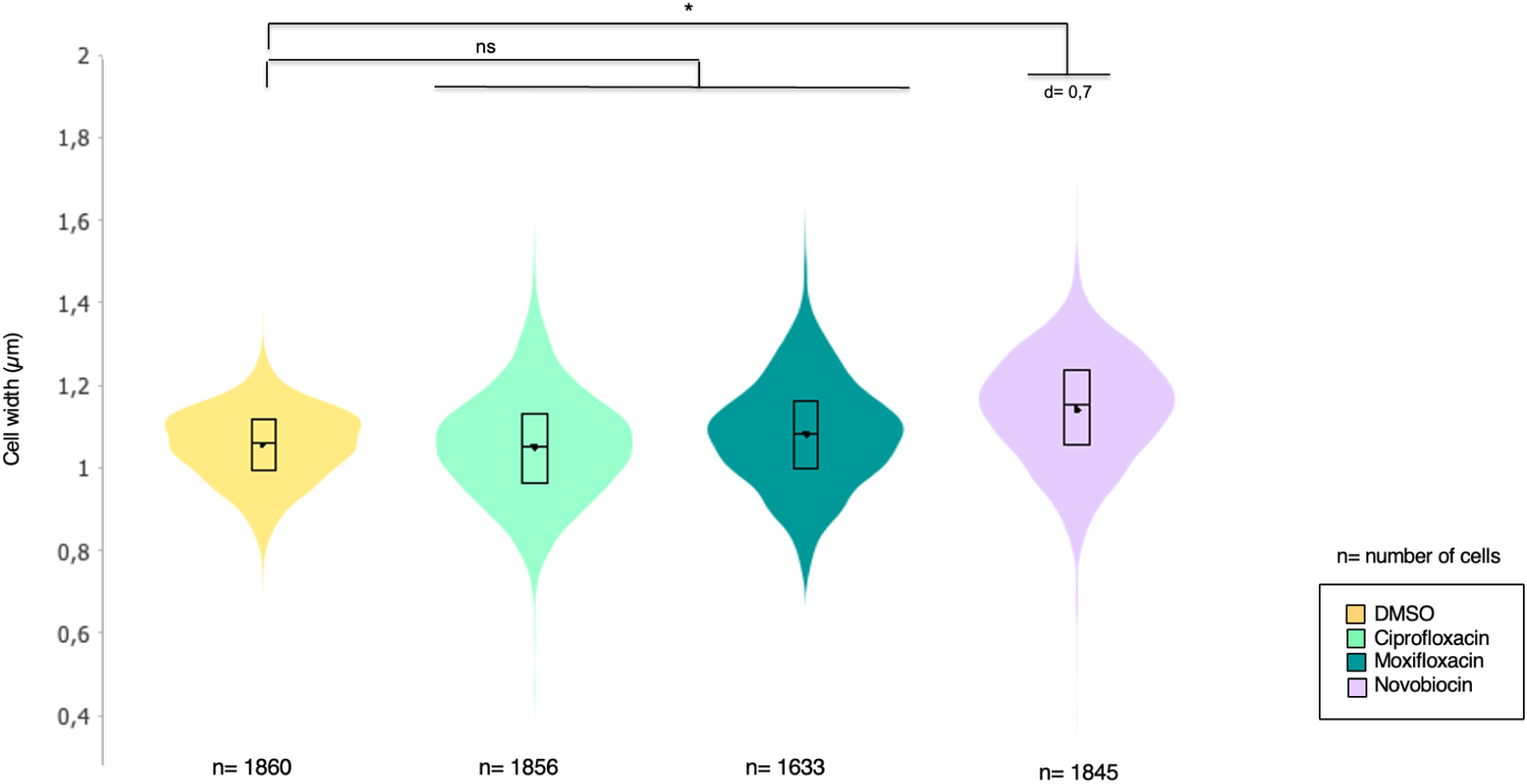
Evaluation of the impact of Ciprofloxacin, Moxifloxacin and Novobiocin on the phenotype of the *Cglu* WT strain ATCC13032. Violin plots illustrating the distribution of cell width for each tested condition from (Figure 4A). (Cohen’s d, from left to right compared to control: (ns, d(Ciprofloxacin) = 0.29, d(Moxifloxacin) = 0.01, *, d(Novobiocin) = 0,7). The box indicates the 25th to 75th percentiles, and whiskers indicate the 95% confidence interval. The mean and median are represented by a dot and a line within the box. The number of cells (n) taken into account for the analysis is indicated below each violin.

**Supplementary Figure 15.**
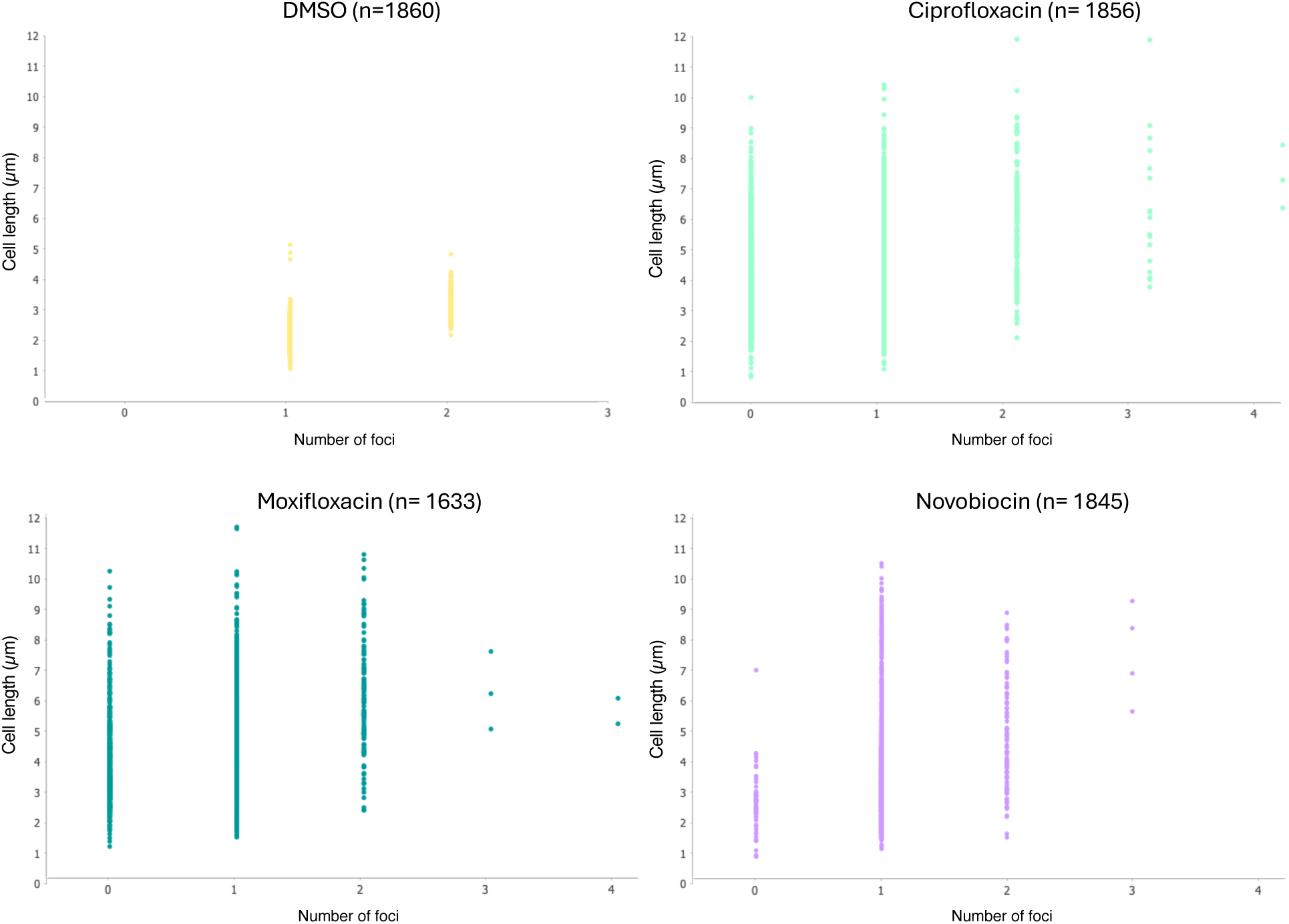
Distribution of cell length according to the number of DNA foci in *Cglu* WT under the effect of different anti-gyrase. Scatter plots show the distribution of cell lengths as a function of the number of fluorescent foci per cell after treatment with DMSO (control), ciprofloxacin, moxifloxacin, or novobiocin. Each dot represents a single cell, n numbers represent the number of cells used in the analyses.

**Supplementary Figure 16.**
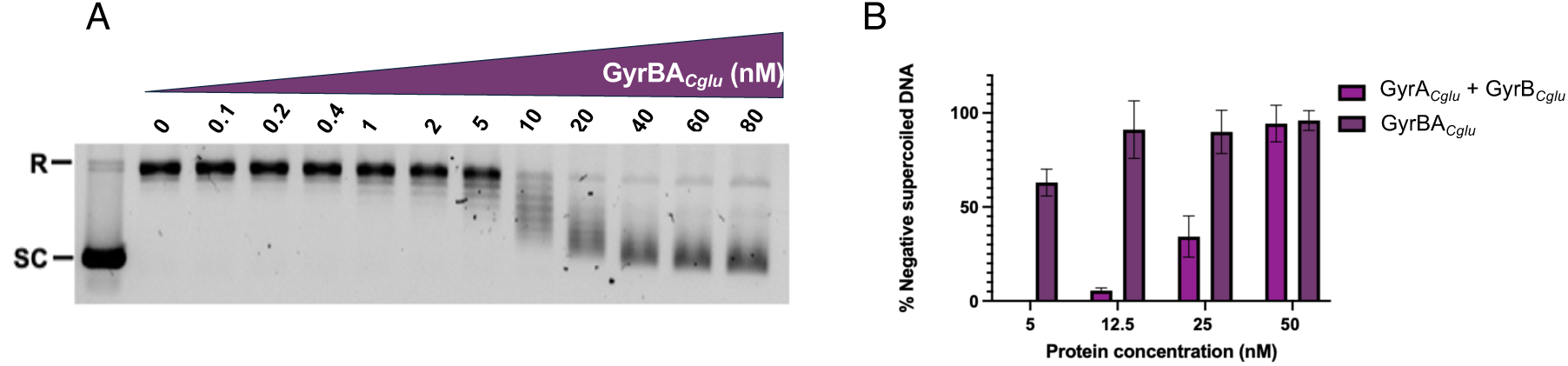
Negative supercoiling activity gyrases. Reactions were carried out in the presence of equimolar quantities of GyrA/GyrB subunits on 300 ng relaxed pBR322 plasmid DNA in the presence of 1 mM ATP. Activity Buffer: 40 mM Tris-HCl (pH 7.5), 25 mM KCl, 6 mM magnesium acetate, 4 mM DTT, 2 mM spermidine, and 0.1 mM EDTA, with the possible addition of potassium glutamate^8^. (A) Fusion GyrBA DNA gyrase of *Cglu*. (B) Comparison of the *Cglu* DNA gyrase negative supercoiling activity for the heterotetramer and the homodimer. R, relaxed DNA; SC, supercoiled DNA;

**Supplementary Figure 18.**
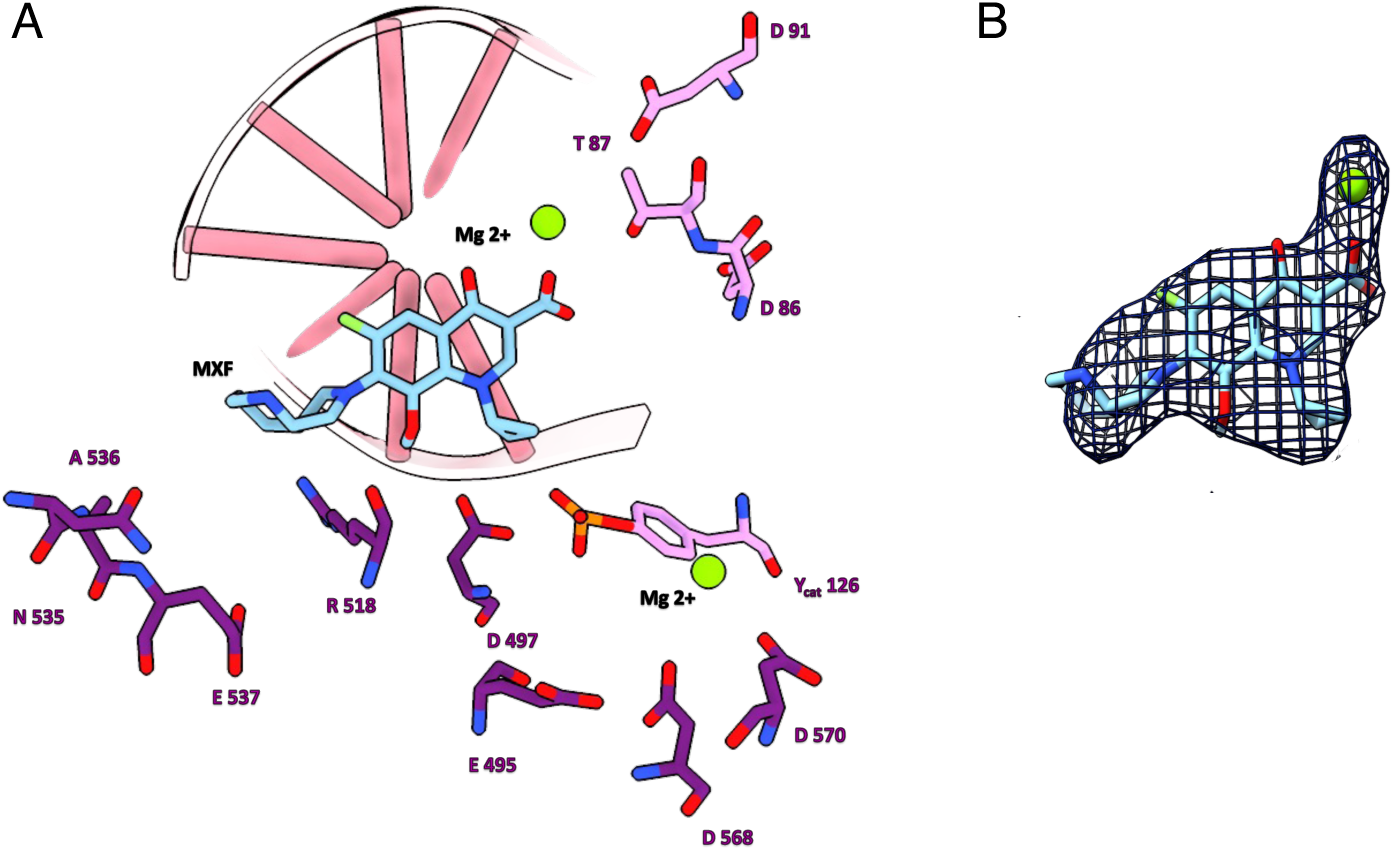
Quinolone binding pocket of *Cglu* gyrase in complex with moxifloxacin. 3D structure of the cryo-EM structure of *Cglu* DNA gyrase. (A) Residues D91, T87, D86 and Y_cat_126 from the GyrA subunit are in light purple and residues A536, N535, E537, R518, D497, E495, D568 and D570 from the GyrB subunit are in dark purple. A molecule of moxifloxacin (MXF, light blue) is intercalated into the cleaved DNA strand near the catalytic tyrosine (Y_cat_126) which is covalently bound to the 5’ end of the cleaved DNA. The first Mg²⁺ (green sphere) is coordinated by the acidic triad E495, D568, and D570 from GyrB and is essential for the cleavage/religation activity of the enzyme. The second Mg²⁺ is important to stabilize MXF via its carboxylate group. Is closed to residues D91, T87, and D86 of GyrA. The ketone group of the quinolone ring is further stabilized by interactions with residues R518 and N535 from GyrB. (B) Moxifloxacin is shown as blue sticks, magnesium ion as green sphere. Cryo-EM map was generated and is shown in dark blue with its accompanying magnesium ion.

**Supplementary Figure 19.**
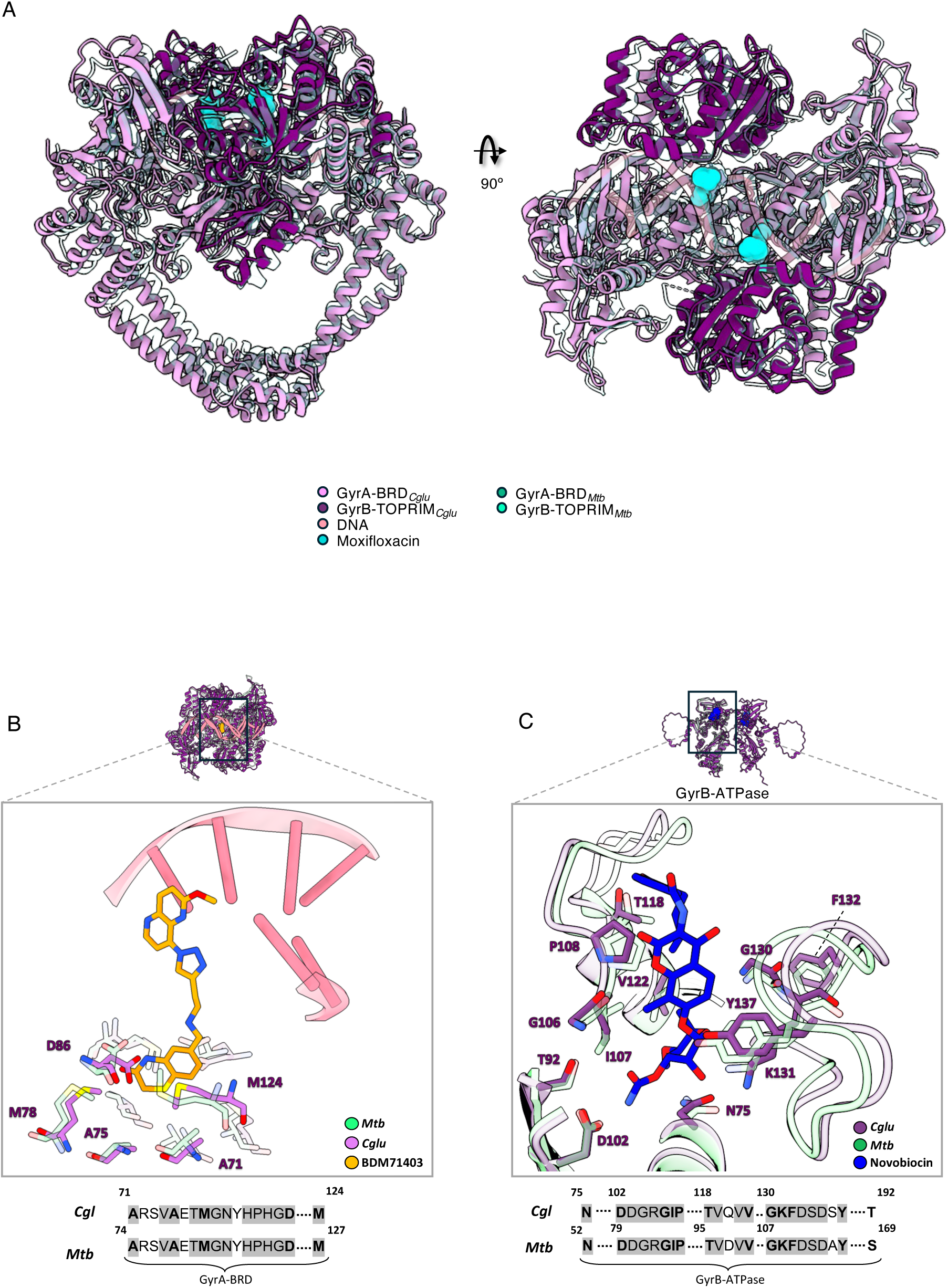
Molecular comparison of DNA gyrase from *Cglu* and *Mtb*: 3D structures comparison and analysis of the inhibitors binding pockets. (A) Superposition of the cleavage core of *Cglu* and *Mtb* gyrases in cartoon representation. Same color code as in Figure 5 panel D for *Cglu*. *Mtb* GyrA and GyrB are shown in transparent light green and dark green respectively. *Mtb* gyrase structure was extracted from PDB 5BS8^9^. (B) **BDM71403-binding pocket:** The cryo-EM structure of *Cglu* DNA gyrase (this work) was superimposed on the cryo-EM structure of *Mtb* gyrase bound to BDM71403 (PDB ID 8S7O). This comparison reveals that the BDM71403 binding pockets of *Cglu* and *Mtb* gyrases are identical. Residues lining the binding sites are shown as stick representations. Color code: *Cglu* in light purple, *Mtb* in green, BDM71403 in orange, and DNA in pink. (C) **Novobiocin-binding pocket:** AF3 model of the *Cglu* ATPase domain was superimposed on the crystallographic structure of *Mtb* gyrase and *Thermus thermophilus* gyrase ATPase domain (PDB ID 3ZKD and 1KIJ, respectively)^11,12^. The ATPase domain from *T. thermophilus* gyrase was resolved in complex with Novobiocin. For better understanding, the structure of *T. thermophilus* is not visible. This comparison reveals that the Novobiocin binding pockets of *Cglu* and *Mtb* gyrases are identical. Residues lining the binding sites are shown as stick representations. Color code: *Cglu* in dark purple, *Mtb* in light green, Novobiocin in blue.

## Supplementary Tables

**Supplementary Table 1.** (A) Physicochemical properties of the GyrA and GyrB subunits of *Cglu* and *Mtb* gyrases. (B) Physicochemical properties of the two domains forming GyrA, the Breakage Reunion Domain (BRD) and the C-terminal domain (CTD) of *Cglu* and *Mtb*. (C) Physicochemical properties of the two domains forming GyrB, the ATPase domain and the TOPoisomerase–PRIMase (TOPRIM) domain from *Cglu* and *Mtb*. (D) Physicochemical properties of the C-tail (the extension of the CTD), the C-loop (insertion in the ATPase domain) and the N-terminal region of the ATPase domain of *Cglu* and *Mtb* gyrases. The residues from the linkers connecting each domain were removed to determine their molecular weights.

| Subunit | GyrA |  | GyrB |  |
| --- | --- | --- | --- | --- |
| Organism | *Cglu* | *Mtb* | *Cglu* | *Mtb* |
| UNIPROT ID | Q8NUC6 | P9WG47 | Q8NUD3 | P9WG45 |
| Molecular weight (Da) | 94 803 | 92 274 | 79 287 | 74 091 |
| Protein sequence identity (%) | 71 |  | 73 |  |
| Protein sequence similarity (%) | 85 |  | 82 |  |

B
| Subunit | GyrA |  |  |  |
| --- | --- | --- | --- | --- |
| Domain | BRD |  | CTD |  |
| Organism | *Cglu* | *Mtb* | *Cglu* | *Mtb* |
| Molecular weight (Da) | 55 075 | 55 319 | 38 102 | 35 287 |
| Protein sequence identity (%) | 75 |  | 64 |  |
| Protein sequence similarity (%) | 97 |  | 96 |  |

C
| Subunit | GyrB |  |  |  |
| --- | --- | --- | --- | --- |
| Domain | ATPase |  | TOPRIM |  |
| Organism | *Cglu* | *Mtb* | *Cglu* | *Mtb* |
| Molecular weight (Da) | 50 552 | 45 368 | 25 003 | 24 976 |
| Protein sequence identity (%) | 68 |  | 83 |  |
| Protein sequence similarity (%) | 97 |  | 100 |  |

D
| Domain | N-ter ATPase |  | C-loop |  | C-tail |  |
| --- | --- | --- | --- | --- | --- | --- |
| Organism | *Cglu* | *Mtb* | *Cglu* | *Mtb* | *Cglu* | *Mtb* |
| Nber of amino acids | 39 | 16 | 45 | 32 | 44 | 22 |
| Acidic amino acids (%) | 10 | 12.5 | 36 | 25 | 29.5 | 27 |
| Basic amino acids (%) | 20.5 | 19 | 11 | 12.5 | 4.5 | 0 |
| Isoelectric point (pI) | 9.9 | 8.2 | 4 | 4.3 | 3.9 | 3.7 |

**Supplementary Table 2.**
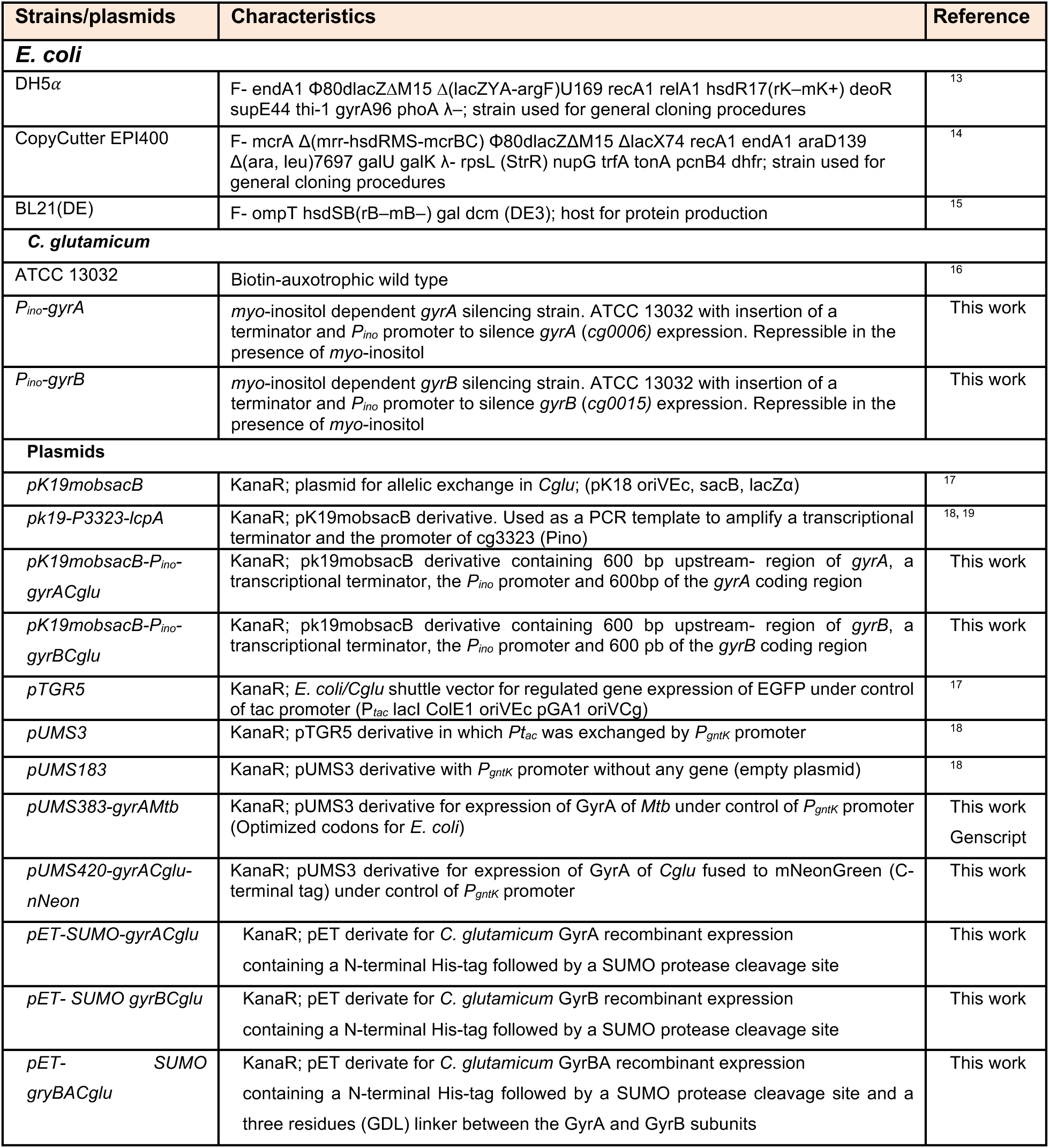
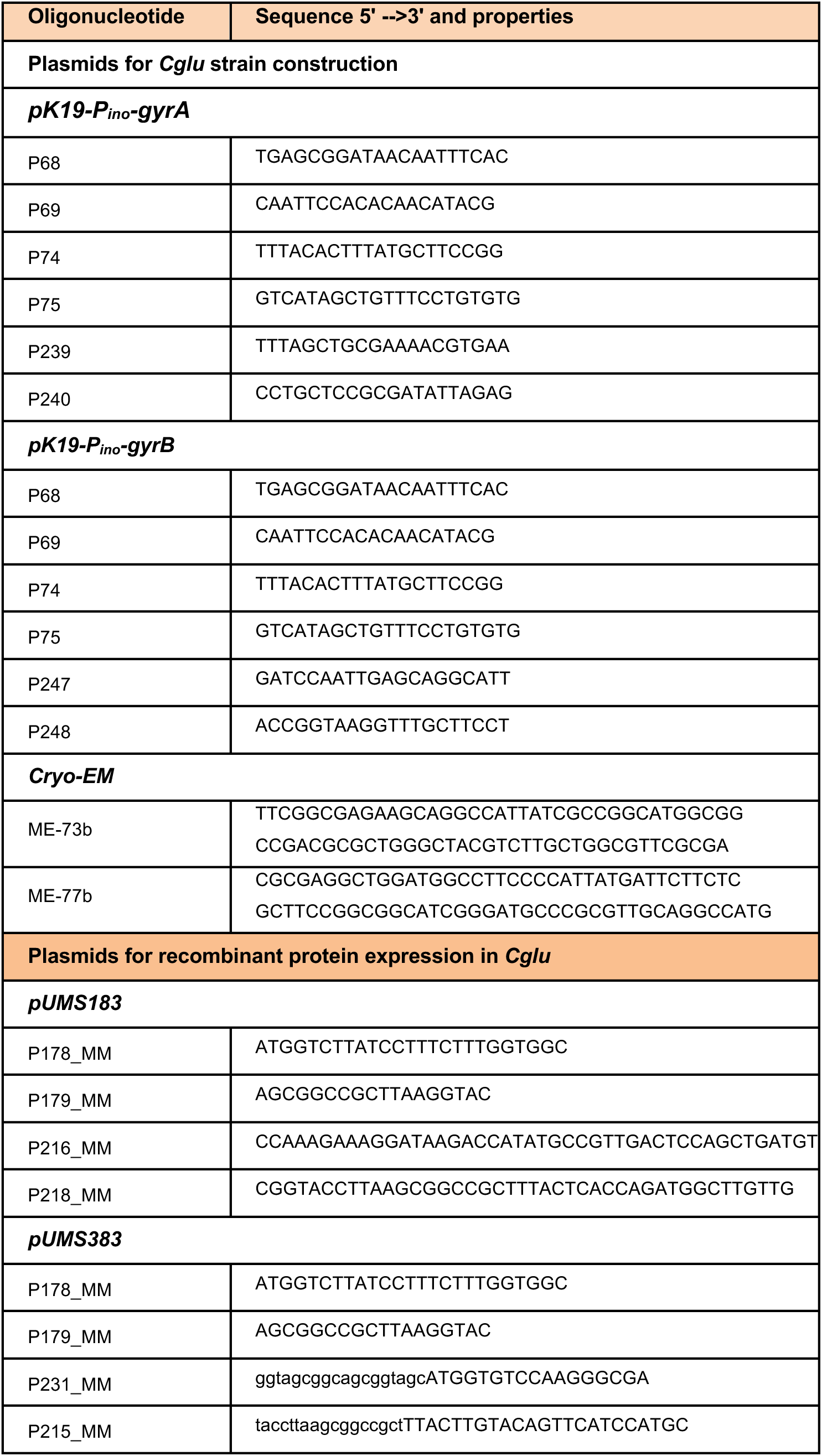
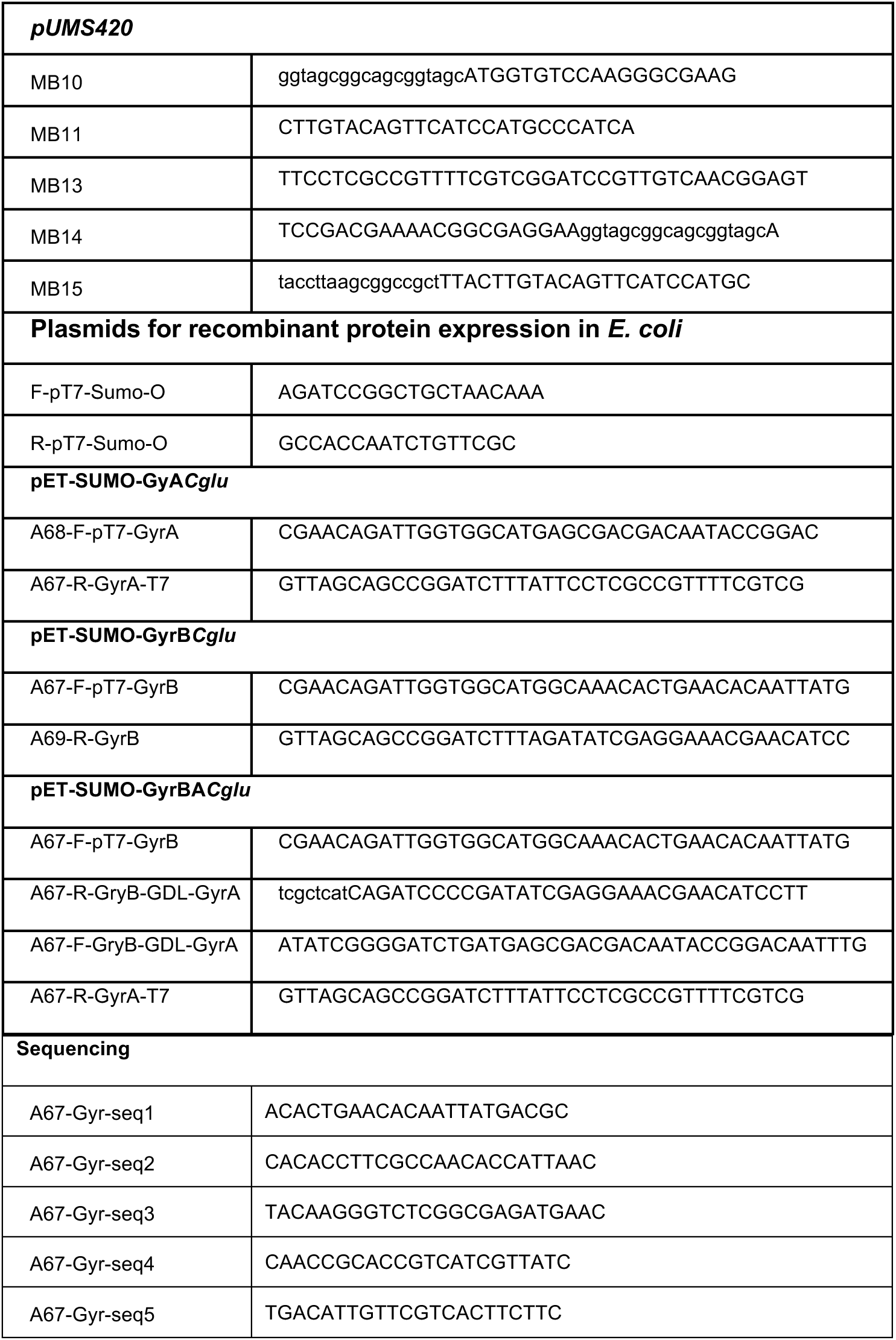
Bacterial strains, plasmids and primers used in this study, with descriptions and references.

## Extended data

**Extended data Figure 1:**
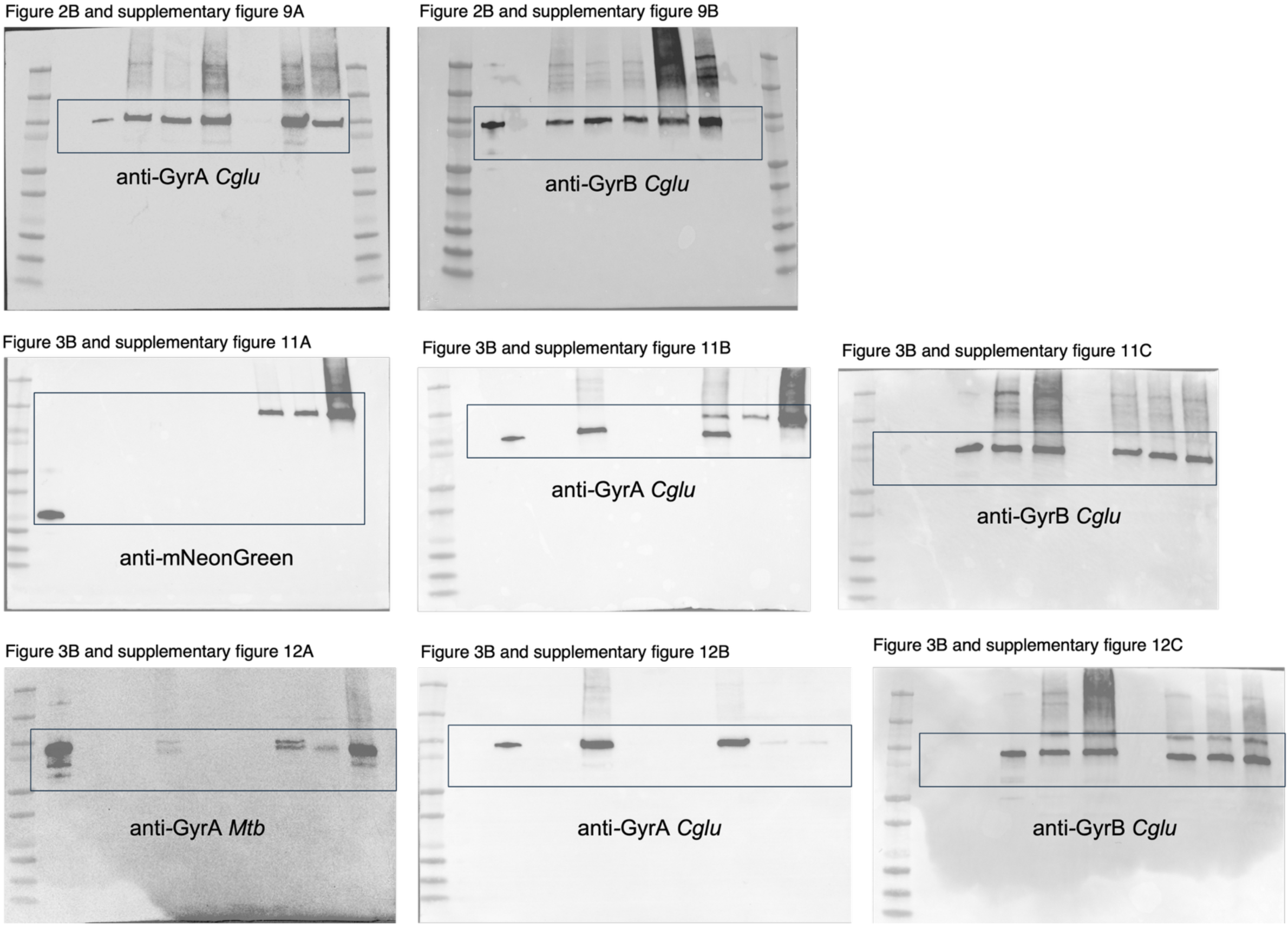
Full uncropped Western Blots of all the analyses shown in this work. The boxes correspond to the crops used in the named figures.

**Extended data Table 1.**
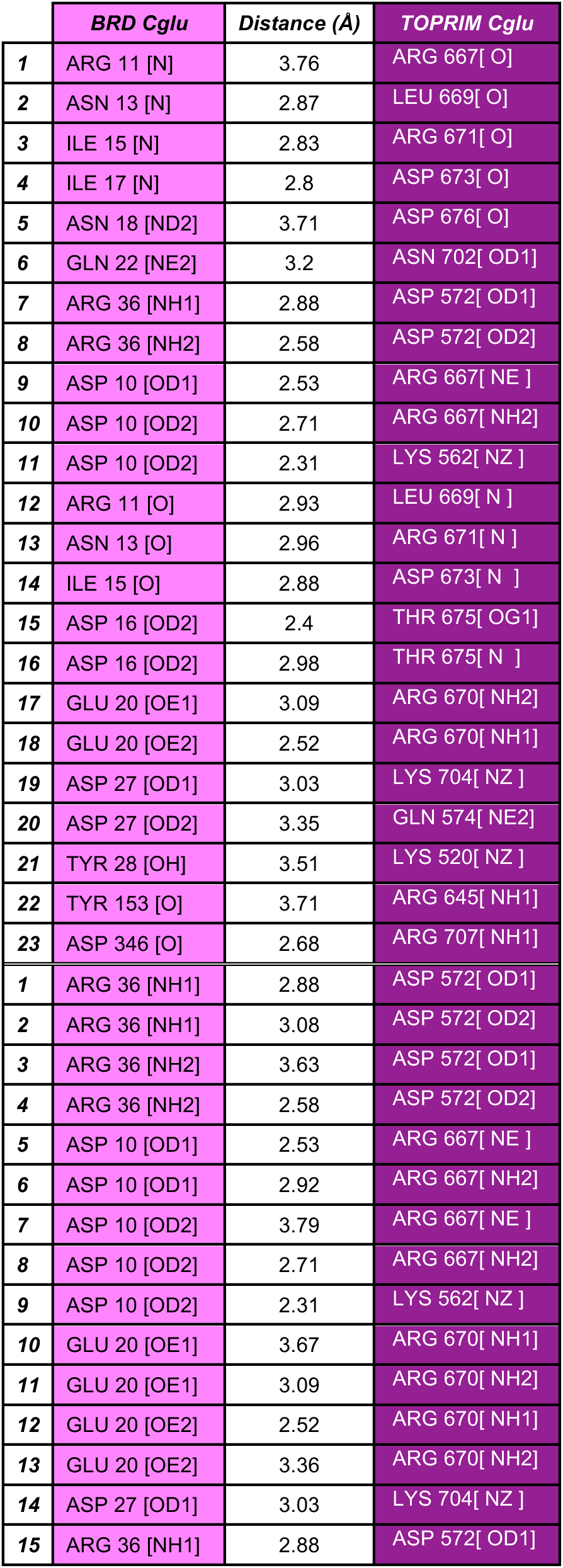
Hydrogen bonds (entries 1 to 23) and Salt bridges (entries 24-43) implicated in GyrB-TOPRIM*_Cglu_*:GyrA-BRD*_Cglu_* dimerisation. Interactions are detected according to a PISA analysis.

|  | *BRD Cglu* | *Distance (Å)* | *TOPRIM Cglu* |
| --- | --- | --- | --- |
| 1 | ARG 11 [N] | 3.76 | ARG 667[ O] |
| 2 | ASN 13 [N] | 2.87 | LEU 669[ O] |
| 3 | ILE 15 [N] | 2.83 | ARG 671[ O] |
| 4 | ILE 17 [N] | 2.8 | ASP 673[ O] |
| 5 | ASN 18 [ND2] | 3.71 | ASP 676[ O] |
| 6 | GLN 22 [NE2] | 3.2 | ASN 702[ OD1] |
| 7 | ARG 36 [NH1] | 2.88 | ASP 572[ OD1] |
| 8 | ARG 36 [NH2] | 2.58 | ASP 572[ OD2] |
| 9 | ASP 10 [OD1] | 2.53 | ARG 667[ NE ] |
| 10 | ASP 10 [OD2] | 2.71 | ARG 667[ NH2] |
| 11 | ASP 10 [OD2] | 2.31 | LYS 562[ NZ ] |
| 12 | ARG 11 [O] | 2.93 | LEU 669[ N ] |
| 13 | ASN 13 [O] | 2.96 | ARG 671[ N ] |
| 14 | ILE 15 [O] | 2.88 | ASP 673[ N ] |
| 15 | ASP 16 [OD2] | 2.4 | THR 675[ OG1] |
| 16 | ASP 16 [OD2] | 2.98 | THR 675[ N ] |
| 17 | GLU 20 [OE1] | 3.09 | ARG 670[ NH2] |
| 18 | GLU 20 [OE2] | 2.52 | ARG 670[ NH1] |
| 19 | ASP 27 [OD1] | 3.03 | LYS 704[ NZ ] |
| 20 | ASP 27 [OD2] | 3.35 | GLN 574[ NE2] |
| 21 | TYR 28 [OH] | 3.51 | LYS 520[ NZ ] |
| 22 | TYR 153 [O] | 3.71 | ARG 645[ NH1] |
| 23 | ASP 346 [O] | 2.68 | ARG 707[ NH1] |
| 1 | ARG 36 [NH1] | 2.88 | ASP 572[ OD1] |
| 2 | ARG 36 [NH1] | 3.08 | ASP 572[ OD2] |
| 3 | ARG 36 [NH2] | 3.63 | ASP 572[ OD1] |
| 4 | ARG 36 [NH2] | 2.58 | ASP 572[ OD2] |
| 5 | ASP 10 [OD1] | 2.53 | ARG 667[ NE ] |
| 6 | ASP 10 [OD1] | 2.92 | ARG 667[ NH2] |
| 7 | ASP 10 [OD2] | 3.79 | ARG 667[ NE ] |
| 8 | ASP 10 [OD2] | 2.71 | ARG 667[ NH2] |
| 9 | ASP 10 [OD2] | 2.31 | LYS 562[ NZ ] |
| 10 | GLU 20 [OE1] | 3.67 | ARG 670[ NH1] |
| 11 | GLU 20 [OE1] | 3.09 | ARG 670[ NH2] |
| 12 | GLU 20 [OE2] | 2.52 | ARG 670[ NH1] |
| 13 | GLU 20 [OE2] | 3.36 | ARG 670[ NH2] |
| 14 | ASP 27 [OD1] | 3.03 | LYS 704[ NZ ] |
| 15 | ARG 36 [NH1] | 2.88 | ASP 572[ OD1] |

**Extended data Table 2.** Hydrogen bonds (entries 1 to 30) and Salt bridges (entries 1-19) implicated in GyrB-TOPRIM*_Cglu_*:GyrA-BRD*_Mtb_* dimerisation. Interactions are detected according to a PISA analysis.

|  | **BRD Mtb** | **Distance (Å)** | **TOPRIM Cglu** |
| --- | --- | --- | --- |
| 1 | ASP 10[ OD1] | 2.28 | LYS 562[ NZ ] |
| 2 | ASP 13[ OD1] | 2.80 | ARG 667[ NE ] |
| 3 | ASP 13[ OD2] | 2.47 | ARG 667[ NH2] |
| 4 | ARG 14[ O] | 2.92 | LEU 669[ N] |
| 5 | GLU 16[ O] | 2.79 | ARG 671[ N] |
| 6 | VAL 18[ O] | 2.83 | ASP 673[ N] |
| 7 | ASP 19[ OD2] | 3.20 | THR 675[ N] |
| 8 | ASP 19[ OD2] | 2.46 | THR 675[ OG1] |
| 9 | GLU 21[ OE1] | 3.00 | ALA 680[ N] |
| 10 | GLU 23[ OE2] | 2.56 | ARG 670[ NH2] |
| 11 | ASP 30[ OD1] | 2.90 | LYS 704[ NZ] |
| 12 | ASP 30[ OD2] | 3.40 | GLN 574[ NE2] |
| 13 | TYR 31[ OH] | 3.35 | LYS 520[ NZ] |
| 14 | TYR 156[ O] | 3.70 | ARG 645[ NH1] |
| 15 | ASP 350[ O] | 2.69 | ARG 707[ NH1] |
| 16 | ASP 350[ OD2] | 3.08 | ARG 707[ NH1] |
| 17 | GLU 16[ N] | 2.87 | LEU 669[ O] |
| 18 | VAL 18[ N] | 2.79 | ARG 671[ O] |
| 19 | ILE 20[ N] | 2.84 | ASP 673[ O] |
| 20 | GLN 25[ NE2] | 3.13 | ASN 702[ OD1] |
| 21 | ARG 39[ NH1] | 2.90 | ASP 572[ OD1] |
| 22 | ARG 39[ NH2] | 2.44 | ASP 572[ OD2] |
| 23 | ARG 159[ NH2] | 3.08 | ASP 641[ OD2] |
| 1 | ASP 10[ OD1] | 2.28 | LYS 562[ NZ] |
| 2 | ASP 10[ OD2] | 3.98 | LYS 562[ NZ] |
| 3 | ASP 13[ OD1] | 3.31 | ARG 667[ NH2] |
| 4 | ASP 13[ OD1] | 2.80 | ARG 667[ NE] |
| 5 | ASP 13[ OD2] | 3.46 | LYS 562[ NZ] |
| 6 | ASP 13[ OD2] | 2.47 | ARG 667[ NH2] |
| 7 | ASP 13[ OD2] | 3.54 | ARG 667[ NE] |
| 8 | GLU 23[ OE1] | 3.44 | ARG 670[ NH1] |
| 9 | GLU 23[ OE2] | 2.85 | ARG 670[ NH1] |
| 10 | GLU 23[ OE2] | 2.56 | ARG 670[ NH2] |
| 11 | ASP 30[ OD1] | 2.90 | LYS 704[ NZ] |
| 12 | ASP 30[ OD2] | 3.97 | LYS 704[ NZ] |
| 13 | ASP 350[ OD2] | 3.08 | ARG 707[ NH1] |
| 14 | ARG 39[ NH1] | 2.90 | ASP 572[ OD1] |
| 15 | ARG 39[ NH1] | 3.06 | ASP 572[ OD2] |
| 16 | ARG 39[ NH2] | 3.67 | ASP 572[ OD1] |
| 17 | ARG 39[ NH2] | 2.44 | ASP 572[ OD2] |
| 18 | ARG 159[ NE] | 3.88 | ASP 641[ OD2] |
| 19 | ARG 159[ NH2] | 3.08 | ASP 641[ OD2] |

**Extended data Table 3.** Hydrogen bonds (entries 1 to 30) and Salt bridges (entries 1-19) implicated in GyrB-TOPRIM*_Mtb_*:GyrA-BRD*_Cglu_* dimerisation. Interactions are detected according to a PISA analysis

|  | **BRD Cglu** | **Distance (Å)** | **TOPRIM Mtb** |
| --- | --- | --- | --- |
| 1 | ARG 11[ NE] | 3.82 | ARG 631[ O] |
| 2 | ARG 11[ NH2] | 2.25 | VAL 630[ O] |
| 3 | ARG 11[ NH2] | 3.55 | SER 629[ O] |
| 4 | ASN 13[ N] | 2.88 | LEU 633[ O] |
| 5 | ILE 15[ N] | 2.86 | GLN 635[ O] |
| 6 | ILE 17[ N] | 2.87 | THR 637[ O] |
| 7 | ASN 18[ ND2] | 3.55 | ASP 640[ O] |
| 8 | GLN 22[ NE2] | 3.49 | ASN 666[ OD1] |
| 9 | TYR 25[ OH] | 2.95 | LEU 651[ O] |
| 10 | ARG 36[ NH1] | 2.84 | ASP 536[ OD1] |
| 11 | ARG 36[ NH2] | 2.56 | ASP 536[ OD2] |
| 12 | ASP 10[ OD1] | 2.91 | ARG 631[ NE] |
| 13 | ASP 10[ OD2] | 2.98 | LYS 526[ NZ] |
| 14 | ASP 10[ OD2] | 2.32 | ARG 631[ NH2] |
| 15 | ARG 11[ O] | 2.93 | LEU 633[ N] |
| 16 | ASN 13[ O] | 2.79 | GLN 635[ N] |
| 17 | ILE 15[ O] | 2.85 | THR 637[ N] |
| 18 | ILE 15[ O] | 3.33 | THR 637[ OG1] |
| 19 | ASP 16[ OD1] | 3.40 | ASP 639[ N] |
| 20 | GLU 20[ OE1] | 3.02 | ARG 634[ NH2] |
| 21 | GLU 20[ OE2] | 2.48 | ARG 634[ NH1] |
| 22 | TYR 25[ OH] | 3.76 | ARG 659[ NH1] |
| 23 | ASP 27[ OD1] | 2.83 | LYS 668[ NZ] |
| 24 | ASP 27[ OD2] | 3.38 | GLN 538[ NE2] |
| 25 | TYR 28[ OH] | 3.55 | LYS 484[ NZ] |
| 26 | TYR 153[ O] | 3.35 | ARG 609[ NH1] |
| 27 | ASP 346[ O] | 2.78 | ARG 671[ NH1] |
| 1 | ARG 36[ NH1] | 2.84 | ASP 536[ OD1] |
| 2 | ARG 36[ NH1] | 3.12 | ASP 536[ OD2] |
| 3 | ARG 36[ NH2] | 3.59 | ASP 536[ OD1] |
| 4 | ARG 36[ NH2] | 2.56 | ASP 536[ OD2] |
| 5 | ASP 10[ OD1] | 2.91 | ARG 631[ NE] |
| 6 | ASP 10[ OD1] | 3.82 | ARG 631[ NH2] |
| 7 | ASP 10[ OD2] | 2.94 | ARG 631[ NE] |
| 8 | ASP 10[ OD2] | 2.98 | LYS 526[ NZ] |
| 9 | ASP 10[ OD2] | 2.32 | ARG 631[ NH2] |
| 10 | GLU 20[ OE1] | 3.63 | ARG 634[ NH1] |
| 11 | GLU 20[ OE1] | 3.02 | ARG 634[ NH2] |
| 12 | GLU 20[ OE2] | 2.48 | ARG 634[ NH1] |
| 13 | GLU 20[ OE2] | 3.34 | ARG 634[ NH2] |
| **14** | ASP 27[ OD1] | 2.83 | LYS 668[ NZ] |
| **15** | ASP 27[ OD2] | 3.86 | LYS 668[ NZ] |

**Extended data Table 4.** Hydrogen bonds (entries 1 to 30) and Salt bridges (entries 1-19) implicated in GyrB-TOPRIM*_Mtb_*:GyrA-BRD*_Mtb_* dimerisation. Interactions are detected according to a PISA analysis.

|  | **BRD Mtb** | **Distance (Å)** | **TOPRIM Mtb** |
| --- | --- | --- | --- |
| 1 | THR 5[ OG1] | 2.39 | MET 474[ O] |
| 2 | ARG 14[ NE] | 3.64 | ARG 631[ O] |
| 3 | ARG 14[ NH2] | 3.79 | ARG 631[ O] |
| 4 | ARG 14[ NH2] | 3.82 | SER 629[ O] |
| 5 | GLU 16[ N] | 2.88 | LEU 633[ O] |
| 6 | VAL 18[ N] | 2.84 | GLN 635[ O] |
| 7 | ILE 20[ N] | 2.90 | THR 637[ O] |
| 8 | GLN 25[ NE2] | 3.17 | ASN 666[ OD1] |
| 9 | TYR 28[ OH] | 2.95 | LEU 651[ O] |
| 10 | ARG 39[ NH1] | 2.89 | ASP 536[ OD1] |
| 11 | ARG 39[ NH2] | 2.56 | ASP 536[ OD2] |
| 12 | ARG 354[ NH1] | 2.70 | ASP 674[ OD2] |
| 13 | ARG 354[ NH2] | 3.88 | ARG 671[ O] |
| 14 | ARG 354[ NH2] | 3.07 | ASP 674[ OD2] |
| 15 | ASP 10[ OD1] | 2.56 | LYS 526[ NZ] |
| 16 | ASP 13[ OD1] | 2.88 | ARG 631[ NE] |
| 17 | ASP 13[ OD2] | 2.84 | LYS 526[ NZ] |
| 18 | ASP 13[ OD2] | 2.41 | ARG 631[ NH2] |
| 19 | ARG 14[ O] | 2.96 | LEU 633[ N] |
| 20 | GLU 16[ O] | 2.78 | GLN 635[ N] |
| 21 | GLU 16[ OE2] | 3.73 | GLN 565[ NE2] |
| 22 | VAL 18[ O] | 2.84 | THR 637[ N] |
| 23 | VAL 18[ O] | 3.36 | THR 637[ OG1] |
| 24 | ASP 19[ OD2] | 3.24 | ASP 639[ N] |
| 25 | GLU 23[ OE1] | 2.97 | ARG 634[ NH2] |
| 26 | GLU 23[ OE2] | 2.48 | ARG 634[ NH1] |
| 27 | ASP 30[ OD1] | 3.06 | LYS 668[ NZ] |
| 28 | ASP 30[ OD2] | 3.50 | GLN 538[ NE2] |
| 29 | TYR 31[ OH] | 3.46 | LYS 484[ NZ] |
| 30 | ASP 350[ O] | 2.89 | ARG 671[ NH1] |
| 1 | ARG 39[ NH1] | 2.89 | ASP 536[ OD1] |
| 2 | ARG 39[ NH1] | 3.17 | ASP 536[ OD2] |
| 3 | ARG 39[ NH2] | 3.57 | ASP 536[ OD1] |
| 4 | ARG 39[ NH2] | 2.56 | ASP 536[ OD2] |
| 5 | ARG 354[ NH1] | 3.10 | ASP 674[ OD1] |
| 6 | ARG 354[ NH1] | 2.70 | ASP 674[ OD2] |
| 7 | ARG 354[ NH2] | 3.17 | ASP 674[ OD1] |
| 8 | ARG 354[ NH2] | 3.07 | ASP 674[ OD2] |
| 9 | ASP 10[ OD1] | 2.56 | LYS 526[ NZ] |
| 10 | ASP 13[ OD1] | 3.38 | ARG 631[ NH2] |
| 11 | ASP 13[ OD1] | 2.88 | ARG 631[ NE] |
| 12 | ASP 13[ OD2] | 2.84 | LYS 526[ NZ] |
| 13 | ASP 13[ OD2] | 2.41 | ARG 631[ NH2] |
| 14 | ASP 13[ OD2] | 3.48 | ARG 631[ NE] |
| 15 | GLU 23[ OE1] | 2.97 | ARG 634[ NH2] |
| 16 | GLU 23[ OE1] | 3.45 | ARG 634[ NH1] |
| 17 | GLU 23[ OE2] | 3.49 | ARG 634[ NH2] |
| 18 | GLU 23[ OE2] | 2.48 | ARG 634[ NH1] |
| 19 | ASP 30[ OD1] | 3.06 | LYS 668[ NZ] |

